# Mitochondrial translation termination, recycling, reinitiation, and rescue for in-frame and out-of-frame contexts

**DOI:** 10.1101/2025.05.03.652009

**Authors:** Taisei Wakigawa, Mari Mito, Qi Fang, Yuzuru Itoh, Yoichi Shinkai, Shintaro Iwasaki

## Abstract

Because mitochondria diverged from a bacterial ancestor during evolution, the mitochondrial protein synthesis system includes both mRNAs and translation factors with unique characteristics. However, the molecular mechanisms underlying translation termination, recycling, and quality control remain unclear. Here, via high-resolution mitochondrial Ribo-Seq and Disome-Seq, we revealed: the specificity of release factors for different kinds of stop codons; the role of mtRF1 in vertebrates, which do not have noncanonical stop codons in their main ORFs; the recycling-coupled translation of internal ORFs; and the rescue of mitoribosomes in the early elongation stage. mtRF1L is a universal release factor that recognizes all stop codons, whereas mtRF1 recognizes only AGA/AGG noncanonical stop codons. Additionally, mtRF1 terminates the translation of out-of-frame ORFs that end with AGA/AGG. We also found that mtRRF and mtIF3 are required for mitoribosome recycling on stop codons and for the reinitiation of internal ORF translation. Mitoribosomes that stall at start codons are major substrates of the rescue factors ICT1, mtRF-R, and mtRES1. Moreover, HEMK1-mediated methylation of release factors enhances the termination reaction on stop codons. Our results provide insights into the mitoribosome dynamics that are associated with the completion of protein synthesis.

## Introduction

The mitochondrial translation system has evolved from a bacterial ancestor, yet it has acquired a set of unique characters that distinguish it from both bacterial and cytosolic translation machineries. These differences include features of mRNAs, such as diverged genetic codes ^1,2^, the polyadenylation-mediated generation of stop codons ^3^, and the polycistronic architecture of open reading frames (ORFs) ^4,5^, as well as features of the translation machinery, such as protein-rich mitochondrial ribosomes (mitoribosomes) ^6^, streamlined initiation factors ^7–9^, and tRNA modifications ^10^. The unique interactions between mRNAs and the translation machinery pose an analytic obstacle in understanding the unusual mechanism of mitochondrial translation.

In addition to the standard stop codons UAG and UAA, two mitochondrial ORFs — *MT-ND6* and *MT-CO1* — end with unconventional stop codons, namely, AGG and AGA, respectively, in humans. The factor(s) that recognizes these noncanonical stop codons has long been under debate ^11,12^. This discussion is strongly associated with two release factors that participate in mitochondrial translation: mtRF1 and mtRF1L ^11,12^ (Figure 1A). mtRF1L has been reported to terminate translation at UAG/UAA canonical stop codons ^13–15^. Additionally, a recent study that used metabolic labeling with ^35^S methionine revealed that mtRF1L facilitates protein synthesis from *MT-ND6*, which ends with AGG, but not *MT-CO1*, which ends with AGA ^16^. The authors concluded that mtRF1, rather than mtRF1L, is responsible for the termination of translation at the AGA stop codon ^16^. In contrast, other reports have shown that mtRF1 is essential for the termination of translation at both the AGG and AGA stop codons ^17,18^. Thus, the stop codon–release factor relationship in mitochondrial protein synthesis remains elusive (Figure 1A). Moreover, the absence of AGA/AGG stop codons in ORFs and the presence of mtRF1 in ∼40% of vertebrates, including mice, further complicate the situation ^11,12^.

**Figure 1.**
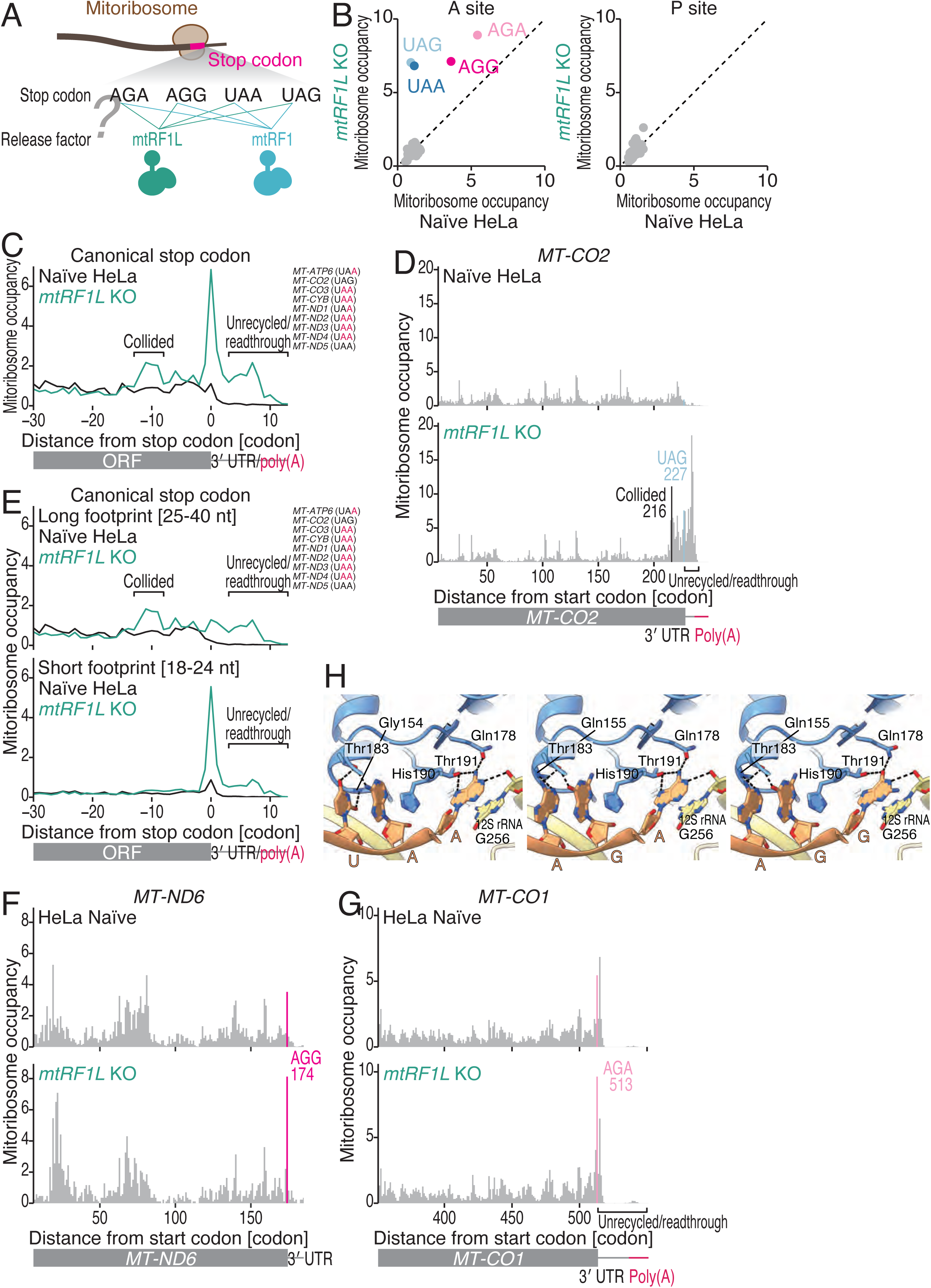
Human mtRF1L recognizes canonical and noncanonical stop codons. (A) Schematic of the questions in this study; the relationship between mitochondrial release factors and stop codon sequences. (B) Comparison of mitoribosome occupancy at the A-site (left) and P-site (right) codons in the indicated cells. (C) Metagene plots of mitoribosome footprints around canonical stop codons in the indicated cells. The A-site position of each read is displayed. The ORFs and stop codon sequences that were used in the analysis are shown. The *MT-ATP8* and *MT-ND4L* ORFs were excluded from the analysis since the 3′ ends of these ORFs overlap with the downstream *MT-ATP6* and *MT-ND4* ORFs in the polycistronic mRNAs. (D, F, and G) Distribution of mitoribosome footprints in naïve and *mtRF1L* KO HeLa cells along the indicated transcripts. The A-site position of each read is displayed. The canonical stop codon and the codon position indicating potential colliding mitoribosomes are highlighted. (E) Same as C but with long footprints (25-40 nt, top) and short footprints (18-24 nt, bottom) separately plotted. (H) Potential interactions of mtRF1L with noncanonical stop codons in the mitoribosome A site. Left: observed interactions of mtRF1L with the UAA codon in the reported cryo-EM structure (PDB ID: 7NQH). Middle and right: modeled structures based on the reported structure. Potential interactions of mtRF1L with the AGA and AGG codons are represented. Hydrogen bonds are shown as dashed lines. See also Figure S1.

Mitochondria also have additional release factors, namely, ICT1 and mtRF-R, which are class 2 release factors. In contrast to class 1 release factors (*e.g.*, mtRF1 and mtRF1L), these proteins lack codon recognition motifs ^11,12^. These factors have been suggested to function in the quality control of translation and the rescue of stalled mitoribosomes ^15,19–25^. However, the natural contexts in which this system functions remain largely unknown.

Translation termination and mitoribosome rescue should be associated with the dissociation of the 39S large mitoribosomal subunit from the 28S small mitoribosomal subunit. This mitoribosome recycling is driven by the mitochondrial recycling factor (mtRRF) in combination with mitochondrial elongation factor G2 (mtEFG2) ^15,26–29^, which is a paralog of the elongation-specific factor mtEFG1. On the basis of what is known about its bacterial homolog, mitochondrial initiation factor 3 (mtIF3) has been suggested to facilitate the recycling process ^30,31^, preventing the reassociation of 39S with 28S on stop codons. Mitoribosome recycling may play a crucial role in balancing protein synthesis from two ORFs in polycistronic mRNAs because the mitoribosomes that function in the translation of upstream ORFs are thought to also function in the translation of downstream ORFs ^32^. Nonetheless, the mechanistic implications of recycling factors in the translation of polycistronic mRNAs have not yet been investigated.

To answer these key questions about mitochondrial protein synthesis, we applied the latest ribosome profiling technology (Ribo-Seq), which is tailored to investigate mitochondrial translation at high resolution ^33^ [mitochondria immunoprecipitation (MitoIP), T7-high resolution original RNA (Thor) Ribo-Seq] to study cells lacking mitochondrial release factors, rescue factors, and recycling factors. Our data revealed the stop codon specificity of mtRF1 (noncanonical AGA/AGG stop codons) and mtRF1L (all four stop codons). Furthermore, by investigating mitoribosome collision via MitoIP-Thor-Disome-Seq ^33^, we found that mtRF1 manages mitoribosome frameshift and recognizes noncanonical codons, especially at the −1 frame, which explains the presence of mtRF1 genes in many vertebrate genomes. Moreover, mtIF3 and mtRRF are required for the efficient dissociation of 39S upon termination and translation from internal ORFs in polycistronic mRNAs, suggesting that 28S slides after the recycling reaction. Moreover, the substantial accumulation of mitoribosomes at start codons when ICT1, mtRF-R, and mtRES1 — a cofactor of mtRF-R— are lacking indicates that mitoribosomes are under surveillance by quality control systems during the early elongation stage. In addition, we showed that Gly-Gly-Gln (GGQ) methylation by HEMK1 promotes the termination reaction. Our data provide insights into the termination, recycling, reinitiation, and rescue processes that are facilitated by mitochondrial translation factors, across mitochonerial transcriptome.

## Results

### mtRF1L recognizes both canonical and noncanonical stop codons

To elucidate the stop codon specificity of mitochondrial release factors (Figure 1A), we performed MitoIP-Thor-Ribo-Seq ^33^. We established HeLa cells in which *mtRF1L* (Figure S1A-D) and *mtRF1* (Figure S2A-C) were deleted via the CRISPR-Cas9 approach.

The *mtRF1L* KO HeLa cells exhibited reduced growth (Figure S1E), even when the medium was supplemented with pyruvate and uridine, which are known to overcome the effects of oxidative phosphorylation (OXPHOS) deficiency ^34^. A similar cell growth phenotype was observed when *mtRF1L* was knocked out in HEK293 cells ^16^ and knocked down in HeLa cells via small interfering RNA (siRNA) ^13^. Furthermore, MitoIP-Thor-Ribo-Seq of *mtRF1L* KO cells (Figure S1F-G) revealed a marked increase in mitoribosome occupancy on standard UAG and UAA stop codons at the A site (Figure 1B). In contrast, mtRF1L depletion had a limited effect on mitoriboome occupancy at the P site (Figure 1B). These findings indicate that mtRF1L terminates translation from ORFs that end with canonical stop codons.

Next, a metagene analysis around the canonical stop codons revealed 2 additional hallmarks (Figure 1C), namely, mitoribosomes 1) formed a line behind mitoribosomes that stalled on UAG/UAA stop codons (Figure 1C, collided), which were located ∼10 codons (*i.e.*, a size of a mitoribosome) upstream of the stop codon, and 2) accumulated at the 3′ untranslated region (UTR) or poly(A) tail (Figure 1C, unrecycled/readthrough). These two features were well illustrated by the distribution of the mitoribosome footprints on *MT-CO2* (Figure 1D), *MT-ND5* (Figure S1H), *MT-ATP8* (Figure S1I), and *MT-ND4L* (Figure S1J). Several phenomena may explain the loss of the 3-nt phase in the 3′ UTR/poly(A) tail reads (Figure S1K): 1) mitoribosomes failed to be recycled, as observed for cytosolic ribosomes ^35–39^; 2) both mitoribosomes reading through stop codons — most likely due to release factor loss ^40,41^ (see the *Limitations* section below for details) — and concomitant frameshifting, which is often associated with ribosome pausing ^42,43^, occurred; and 3) the combination of 1) and 2).

During our analysis, we noticed that the accumulation of footprints on the standard stop codons predominantly originated from 18–24 nt short reads (Figures 1E and S1L). These short footprints represented a minor but significant population of the entire mitoribosome footprints (Figure S1F). Similar short mitoribosome footprints were previously reported ^44^. Previous studies have shown that cytoribosome footprints that are generated by ribosomes free from A-site tRNAs or eRF1, which is a release factor that participates in cytosolic translation, lead to 3′ trimming by RNase and thus to the generation of shorter footprints ^45,46^. Our short mitoribosome footprints were trimmed from the 3′ ends with the same 15-nt offset relative to the A-site as the offset that was observed in long 32-nt footprints (Figure S1L-M). Given the reduced availability of mitochondrial release factors to stop codons at the A site, these short footprints represented A site-free mitoribosomes. We noted that the mitoribosomes that trailed paused mitoribosomes at canonical stop codons in *mtRF1L* KO cells were associated with A-site mitochondrial tRNA (mt-tRNA) since long footprints were dominant (Figure 1E). Notably, in addition to the canonical UAG/UAA stop codons, mtRF1L depletion resulted in the accumulation of footprints on the noncanonical AGG/AGA stop codons (Figure 1B and 1F-G). These data showed that mtRF1L is a universal release factor that functions at both canonical and noncanonical stop codons. Consistent with this idea, on-gel mitochondrial-specific fluorescent noncanonical amino acid tagging (on-gel mito-FUNCAT) ^47,48^ revealed a global reduction in mitochondrial translation, including the translation of *MT-CO1* and *MT-ND6,* which end with noncanonical stop codons (Figure S1N-O). Western blotting also revealed that reduced protein level of MT-CO1 (Figure S1P-Q).

To determine whether the recognition of the noncanonical stop codons by mtRF1L is structurally reasonable, we modeled the structures of the interactions of mtRF1L with AGA and AGG in the mitoribosome A site on the basis of the reported cryo-EM structure of a complex containing a mitoribosome and mtRF1L interacting with the UAA codon (PDB ID: 7NQH) ^15^. In this cryo-EM structure, mtRF1L has six hydrogen bonds and one π-stacking interaction with the UAA nucleotides, in which Gly154 (main chain), Thr183, Thr191, Gln178, and His190 (π-stacking) participate (Figure 1H, left). However, the modeled structures suggested the formation of six and five hydrogen bonds with the AGG and AGG codons, respectively, as well as one π-stacking interaction (Figure 1H, middle and right). The mtRF1L residues that are involved in this interaction are mostly the same except for Gly154, while Gly155 also interacts with the AGA and AGG nucleotides. These findings suggest that mtRF1L can interact with the AGA and AGG nucleotides in the mitoribosome A site and thus recognize these noncanonical stop codons.

### mtRF1 specifically recognizes noncanonical stop codons

In contrast to mtRF1L-depleted cells, *mtRF1* KO HeLa cells were relatively healthy and did not exhibit drastic growth inhibition (Figure S2D), and these results were consistent with results of mtRF1-deficient HEK293 cells ^16,17^. MitoIP-Thor-Ribo-Seq of *mtRF1* KO cells (Figure S2E-F) revealed notable mitoribosome stalling on both the AGG codon of *MT-ND6* and the AGA codon of *MT-CO1* (Figure 2A-C). Moreover, we observed the formation of mito-disomes on the AGG stop codon (Figure 2B). A similar but weaker enrichment of colliding mitoribosome was observed upstream of the AGA codon (Figure 2C). Furthermore, long footprints were predominantly observed, which suggested that trailing mitoribosomes collided after mt-tRNA delivery to the A site (Figure S2G-H).

**Figure 2.**
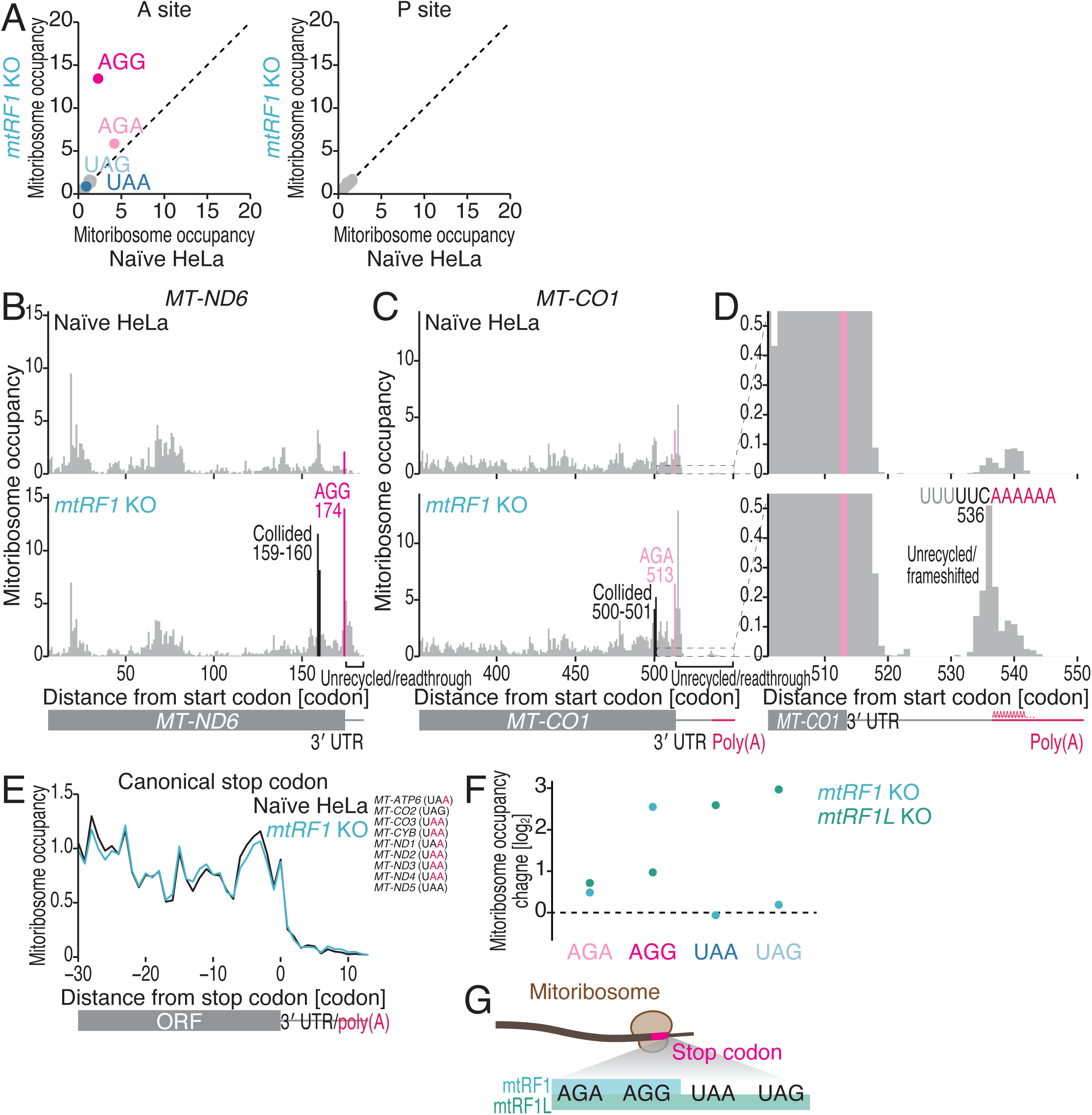
Human mtRF1 recognizes noncanonical stop codons. (A) Comparison of mitoribosome occupancy at the A-site (left) and P-site (right) codons in the indicated cells. (B-D) Distribution of mitoribosome footprints in naïve and *mtRF1* KO HeLa cells along the indicated transcripts. The A-site position of each read is displayed. The noncanonical stop codons and the codon positions indicating potential colliding mitoribosomes are highlighted. D shows a magnified view of C downstream of the stop codon. (E) Metagene plots of mitoribosome footprints around canonical stop codons in the indicated cells. The A-site position of each read is displayed. The ORFs and stop codon sequences that were used in the analysis are shown. The *MT-ATP8* and *MT-ND4L* ORFs were excluded from the analysis since the 3′ ends of these ORFs overlap with the downstream *MT-ATP6* and *MT-ND4* ORFs in the polycistronic mRNAs. (F) Changes in mitoribosome occupancy at stop codons in the indicated cells. (G) Schematic of the relationship between mitochondrial release factors and stop codon sequences. See also Figure S2.

In addition, we observed that mitoribosomes accumulated downstream of the AGG and AGA codons (Figure 2B-C), reaching the boundary of the 3′ UTR and poly(A) tail in the case of *MT-CO1* (Figure 2D). These mitoribosomes were not located at a specific frame, as shown by the absence of 3-nt periodicity (Figure S2I). These features phenocopied the features observed in canonical stop codons in *mtRF1L* KO cells (Figures 1 and S1).

In sharp contrast to mtRF1L depletion, mtRF1 deficiency did not affect translation events at the canonical UAG/UAA stop codons; we did not observe mitoribosome stalling on those codons, mitoribosome collision, or 3′ UTR/poly(A) tail reads (Figure 2A and 2E). On-gel mito-FUNCAT of *mtRF1* KO cells also confirmed this conclusion, revealing a profound reduction in the levels of newly synthesized MT-CO1, which possesses an AGA stop codon (Figure S2J-K). Reduced steady-state level of MT-CO1 protein was also evident by Western blot (Figure S1P-Q). Overall, we concluded that mtRF1 is a release factor that specifically recognizes noncanonical stop codons but not canonical stop codons.

A comparison of mitoribosome stalling at stop codons highlighted the effects of mtRF1 and mtRF1L (Figure 2F-G). Notably, the impacts of both release factors on the AGA codon were comparable, whereas mtRF1 contributed more strongly to translation termination at the AGG codon. Thus, redundant recognition of noncanonical stop codons by two release factors ensures the proper termination of translation in mitochondria.

### The human and mouse mtRF1 proteins rescue mitoribosomes translating out-of-frame ORFs

The high specificity of mtRF1 for AGG/AGA stop codons led us to investigate the function of mouse mtRF1 since none of the ORFs that are encoded in the mouse mitochondrial genome rely on AGG/AGA stop codons (Figure 3A) ^5,11,12^. To explore the role of mtRF1 in mice, we knocked out *mtRF1* in mouse C2C12 cells by the CRISPR-Cas9 method (Figure S3A-C). mtRF1 depletion in C2C12 cells did not impact cell growth (Figure S3D) or global mitochondrial protein synthesis (Figure S3E-F). MitoIP-Thor-Ribo-Seq of mouse *mtRF1* KO cells (Figure S3G-H) revealed a limited effect on mitoribosome occupancy across different types of codons (Figure S3I). As expected, mtRF1 depletion in mice did not lead to the accumulation of footprints on canonical stop codons (Figure S3I-J).

**Figure 3.**
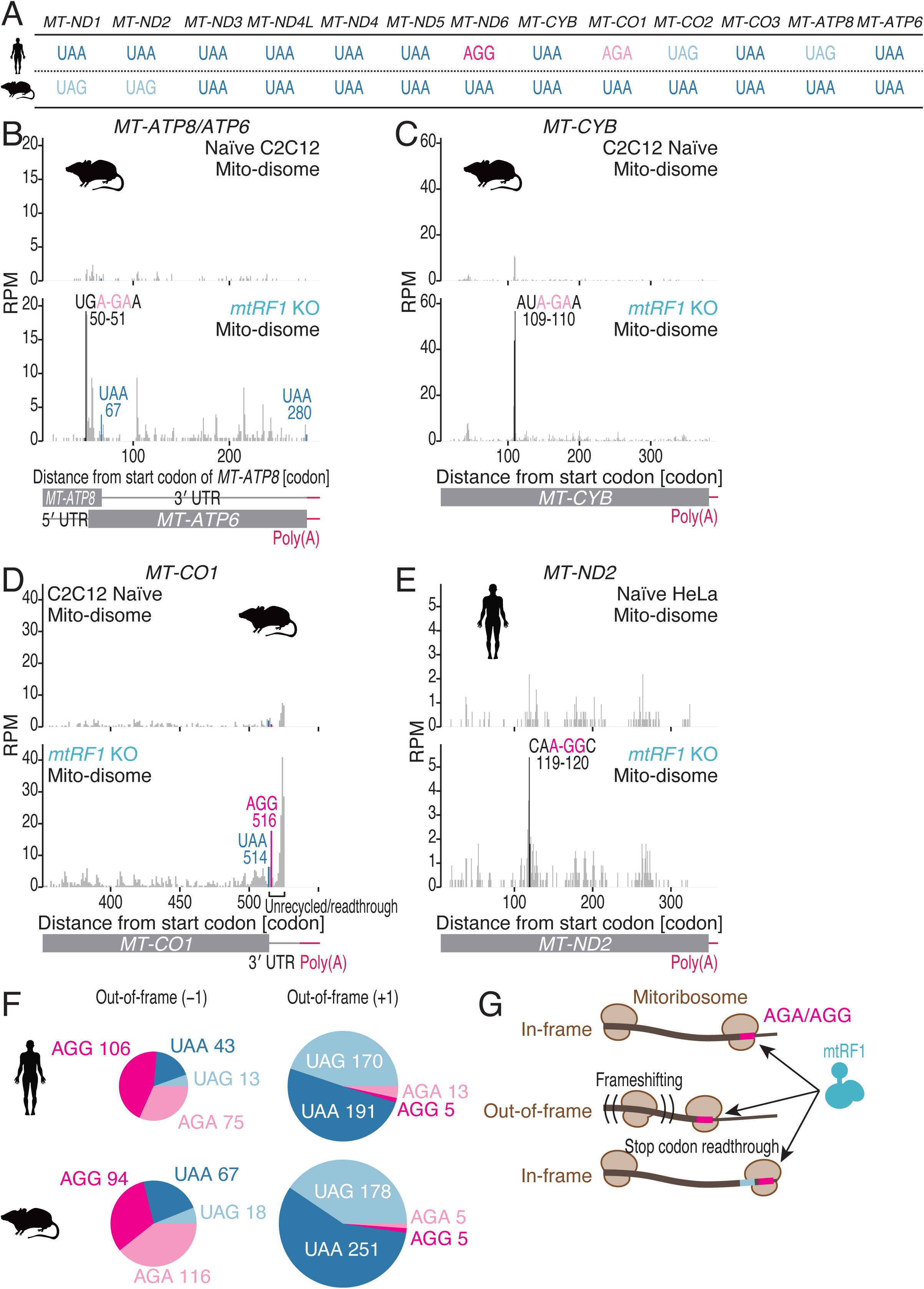
Mouse mtRF1 manages translation termination at out-of-frame ORFs. (A) Schematic of the stop codon sequences in each ORF in humans and mice. (B-D) Distribution of mito-disome footprints in naïve and *mtRF1* KO C2C12 cells along the indicated transcripts. The A-site position (for the leading mitoribosome) of each read is displayed. The noncanonical stop codons and the codon positions with strong peaks are highlighted. RPM, reads per million mapped reads. (E) Distribution of mito-disome footprints in naïve and *mtRF1* KO HeLa cells along the indicated transcripts. The A-site position (for the leading mitoribosome) of each read is displayed. The codon positions with strong peaks are highlighted. RPM, reads per million mapped reads. (F) Pie chart of the numbers of UAA, UAG, AGA, and AGG stop codons in the −1 (left) and +1 (right) frames in human (top) and mouse (bottom) mitochondrial mRNAs. (G) Schematic of the RNA contexts targeted by mtRF1. See also Figures S3-5.

The difficulty in observing the translational outcome of mtRF1 depletion in mouse cells led us to perform a more sensitive assay to survey mitoribosome stalling associated with termination deficiency (Figures 1-2). Thus, we conducted MitoIP-Thor-Disome-Seq, which measures the longer footprints generated by colliding mitoribosomes (or mito-disomes) ^33^. Even with this technique (Figure S3K), we did not observe the accumulation of mito-disomes on canonical stop codons (Figure S3L-M). However, we found two codon positions (in the ORFs of *MT-ATP8* and *MT-CYB*) with a high accumulation of mito-disomes (Figure 3B-C). Importantly, those sites contained the AGA sequence at the −1 frame (Figure 3B-C). Notably, we could not observe these pause events in the mito-monosome footprints (*i.e.*, MitoIP-Thor-Ribo-Seq), possibly due to the difficulty in dissecting the regularly elongating and stalled mitoribosomes (Figure S3N-O).

These data suggest that mitoribosomes may have a significant chance of translating “out-of-frame” ORFs in mRNAs. A recent study that involved investigating secondary structures in human cells revealed that a stem loop in the *MT-ATP8*/*MT-ATP6* mRNA serves as a roadblock for mitoribosome traversal and induces a −1 frameshift that is associated with slippery sequences (Figure S3P, top) ^49^. Translation is stopped by the UAA codon to generate a truncated form of the MT-ATP8 protein (Figure S3P, top) ^49^. Analogous to the human situation, a stem loop fold was predicted at a similar position on the *MT-ATP8*/*MT-ATP6* mRNA (by MXfold2 ^50^) (Figure S3P, bottom). However, the mouse *MT-ATP8* ORF does not possess the corresponding UAA stop codon in the −1 frame, but rather, the AGA stop codon is present downstream (Figure S3P, bottom). Indeed, this noncanonical stop codon in the −1 frame resulted in the formation of mito-disomes when mtRF1 was deleted (Figure 3B). These data suggest that −1-frameshifts may occur on the *MT-ATP8*/*MT-ATP6* mRNA in mouse cells, as suggested in human cells, but this frameshift may generate a slightly different truncated protein.

In addition to frameshifts, mouse mtRF1 may terminate translation after stop codon readthrough. We noticed that a mito-disome formed on the 3′ UTR of *MT-CO1* at the AGA codon, which is located two-codon downstream of the UAA stop codon (Figures 3D and S3Q). This suggested the presence of a mitoribosome subpopulation that may constantly read through stop codons ^33^.

To investigate whether the role of mtRF1 in the context of out-of-frame ORFs is restricted to mice, we performed MitoIP-Thor-Disome-Seq on mtRF1-depleted human HeLa cells (Figure S4A). As expected based on the mito-monosome footprints (Figure 2A and 2C), we observed that mito-disomes were enriched at the AGA noncanonical stop codon (Figure S4B-C). Again, mito-disome pause sites were found in the middle of the ORFs, and those sites were associated with out-of-frame noncanonical stop codons (Figures 3E and S4D-E).

Taken together, our data revealed that mtRF1 plays a critical role in completing protein synthesis from out-of-frame ORFs with the AGG/AGA stop codons and in-frame 3′ UTR extensions (Figure S4F). Consistent with our survey, the human and mouse mitochondrial genomes have greater rates of AGG/AGA occurrence in ORFs with −1 frameshifts than in ORFs with +1 frameshifts (Figure 3F). Generally, the −1 frameshift is more common than the +1 frameshift ^42,43^. mtRF1 may serve as a factor that rescues programmed and/or spontaneous −1 frameshifting during mitochondrial translation (Figure 3G) and therefore is maintained in vertebrates even though the main mitochondrial ORFs do not end with the AGG/AGA codons.

### The use of mito-disomes for stop codon annotation

Given the high sensitivity of mito-disome footprints for detecting hidden noncanonical stop codons, we reasoned that MitoIP-Thor-Disome-Seq of *mtRF1L* KO HeLa cells may be useful for further investigating stop codons in mitochondrial mRNAs that have not yet been annotated. Mito-disomes generated upon mtRF1L depletion (Figure S5A) were found in annotated canonical stop codons (Figure S5B-E). As shown in the mito-monosome footprints (Figure 2F), mito-disome accumulation also recapitulated the similar specificity of the release factors for different types of codons (Figure S5G). Moreover, we observed mito-disomes reaching the 3′ UTR/poly(A) tail (Figure S5C), as mitoribosomes that were not recycled and/or mitoribosomes that were associated with stop codon readthrough/frameshifting (as shown in Figure 1C) collided with trailing mitoribosomes on ORFs.

Our mito-disome footprint data revealed stop codons in short ORFs found in the 3′ UTR. Previous studies have shown that the 3′ UTR of *MT-ND5* has short ORFs [*MT-ND5* downstream ORF (*MT-ND5-dORF*)] ^33,51^. Although the straightforward interpretation of the nucleotide sequence indicated that the dORF contains only 4 amino acids followed by the UAA stop codon, careful data analysis suggested that translation may be terminated at the next in-frame stop codon, namely, the UAG codon, generating a peptide 20 amino acids in length ^51^. Our MitoIP-Thor-Disome-Seq data under conditions of mtRF1L depletion revealed a high accumulation of reads at the second UAG stop codon of *MT-ND5-dORF* (Figure S5H), indicating that a substantial fraction of the mitoribosomes can read through the first stop codon to synthesize longer peptides.

### mtRRF and mtIF3 facilitate mitoribosome recycling and internal ORF translation

Our analysis of mito-monosome footprints in *mtRF1L* KO HeLa cells revealed that termination defects in upstream ORFs in polycistronic mRNAs affected the mitoribosome load on the downstream ORFs; that is, the mitoribosome densities in the internal ORFs *MT-ATP6* and *MT-ND4* were reduced (Figure S1R) when mitoribosomes accumulated at the stop codons of the upstream ORFs *MT-ATP8* and *MT-ND4L*, respectively (Figure S1I-J). This observation was consistent with the idea that upstream ORF translation is required for downstream ORF translation ^32^, which is termed the reinitiation model. One probable scenario is that the mitoribosome that functions in the translation of the upstream ORF is recycled and reused in the translation of the downstream ORF.

To test this hypothesis, we knocked out mitochondrial ribosome recycling factor (mtRRF) in HeLa cells (Figure S6A-C). Depletion of this protein resulted in slower cell growth (Figure S6D), reduced global mitochondrial translation (Figure S6E-F), and decreased the corresponding proteins (Figure S6G-H); similar results were observed when mtRRF was knocked down in HeLa cells by siRNA ^27^. MitoIP-Thor-Ribo-Seq of this cell line (Figure S6I-J) revealed the accumulation of mitoribosomes on both canonical and noncanonical stop codons (Figure 4A-D) due to defects in the dissociation of 39S from 28S after the termination reaction. Consistent with this, we observed A site-free mitoribosomes on the stop codon (Figure 4F) and sliding unrecycled mitoribosomes on the 3′ UTR/poly(A) tail (Figures 4E-F and S6K-L), which did not exhibit 3-nt periodicity (Figure S6M). Ultimately, we observed reduced translation from downstream the *MT-ATP6* and *MT-ND4* ORFs (Figures 4G and S6N-O), indicating the necessity of 39S dissociation after termination of the translation of the first ORF in order for the second ORF to be translated in polycistronic mRNAs.

**Figure 4.**
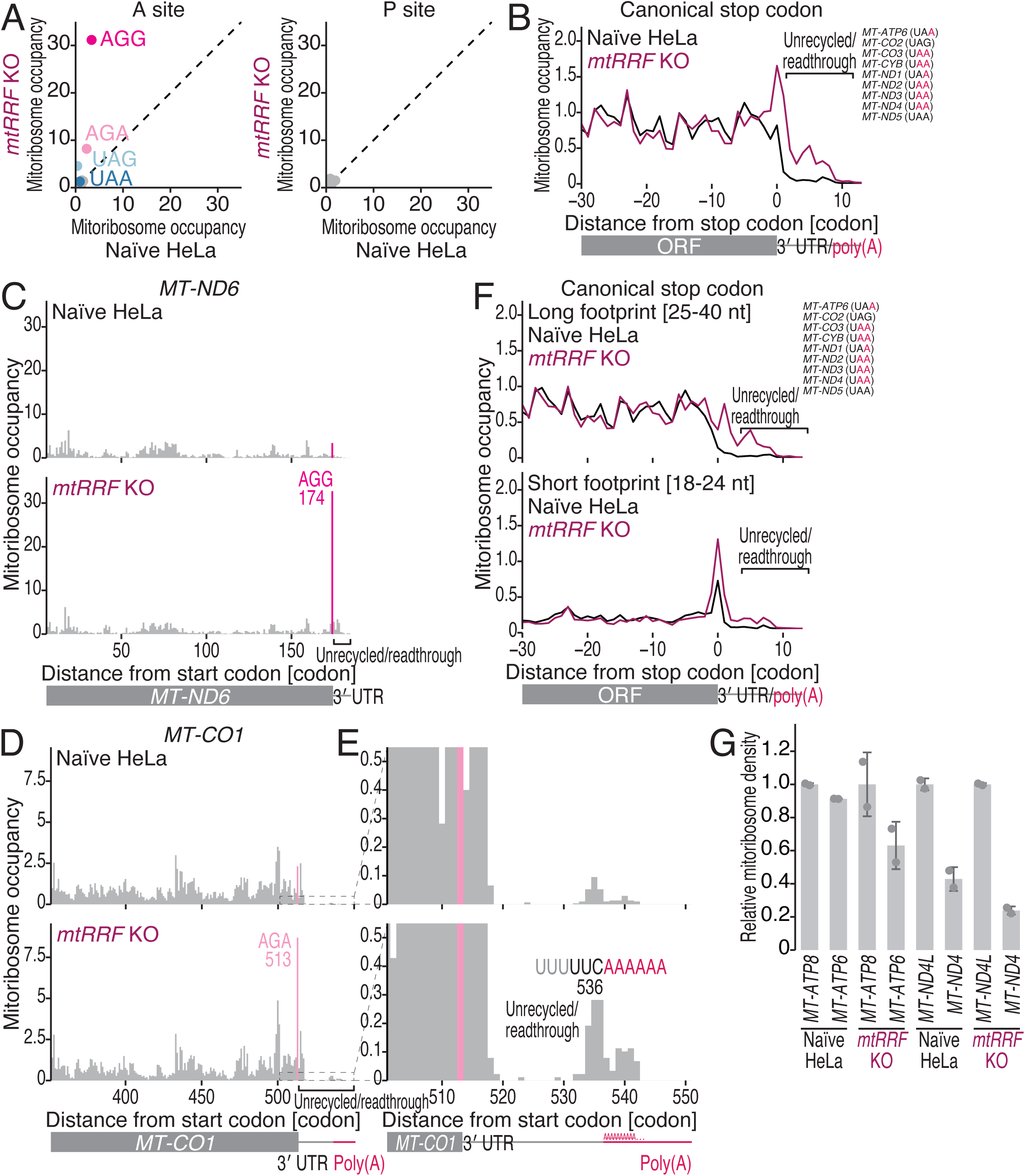
mtRRF is required for mitoribosome recycling and internal ORF translation. (A) Comparison of mitoribosome occupancy at the A-site (left) and P-site (right) codons in the indicated cells. (B) Metagene plots of mitoribosome footprints around canonical stop codons in the indicated cells. The A-site position of each read is displayed. The ORFs and stop codon sequences that were used in the analysis are shown. The *MT-ATP8* and *MT-ND4L* ORFs were excluded from the analysis since the 3′ ends of these ORFs overlap with the downstream *MT-ATP6* and *MT-ND4* ORFs in the polycistronic mRNAs. (C-E) Distribution of mitoribosome footprints in naïve and *mtRRF* KO HeLa cells along the indicated transcripts. The A-site position of each read is displayed. The noncanonical stop codons are highlighted. E represents the magnified view of D downstream of the stop codon. (F) The same as B but with long footprints (25-40 nt, top) and short footprints (18-24 nt, bottom) separately plotted. (G) Relative mitoribosome density on the ORFs in polycistronic mRNAs in the indicated cells. The data are normalized to the upstream ORFs (*MT-ATP8* and *MT-ND4L*) in the polycistronic mRNAs. The means (bars), s.d.s (errors), and individual replicates (n = 2, points) are shown. See also Figure S6.

Recently, mtIF3 was reported to be required for downstream (but not upstream) ORF translation ^33,52,53^. This led us to hypothesize that mtIF3 may facilitate mitoribosome recycling after the termination reaction. Consistent with this expectation, MitoIP-Thor-Ribo-Seq of *mtIF3* KO HEK293 cells (Figure S7A-B), which were generated in an earlier study ^33^, revealed predominant mitoribosome stalling at both canonical and noncanonical stop codons (Figure 5A-D). The accumulation of short footprints represented unrecycled mitoribosome fractions with free A sites (Figure 5E). Considering that mtIF3 has the ability to prevent subunit reassociation ^30,31^, the depletion of mtIF3 may lead to an equilibrium where more 39S subunits do not dissociated from 28S subunits at the stop codon. As observed when *mtRRF* was knocked out, mtIF3 depletion reduced mitoribosome accumulation on the internal *MT-ATP6* and *MT-ND4* ORFs (Figure 5F).

**Figure 5.**
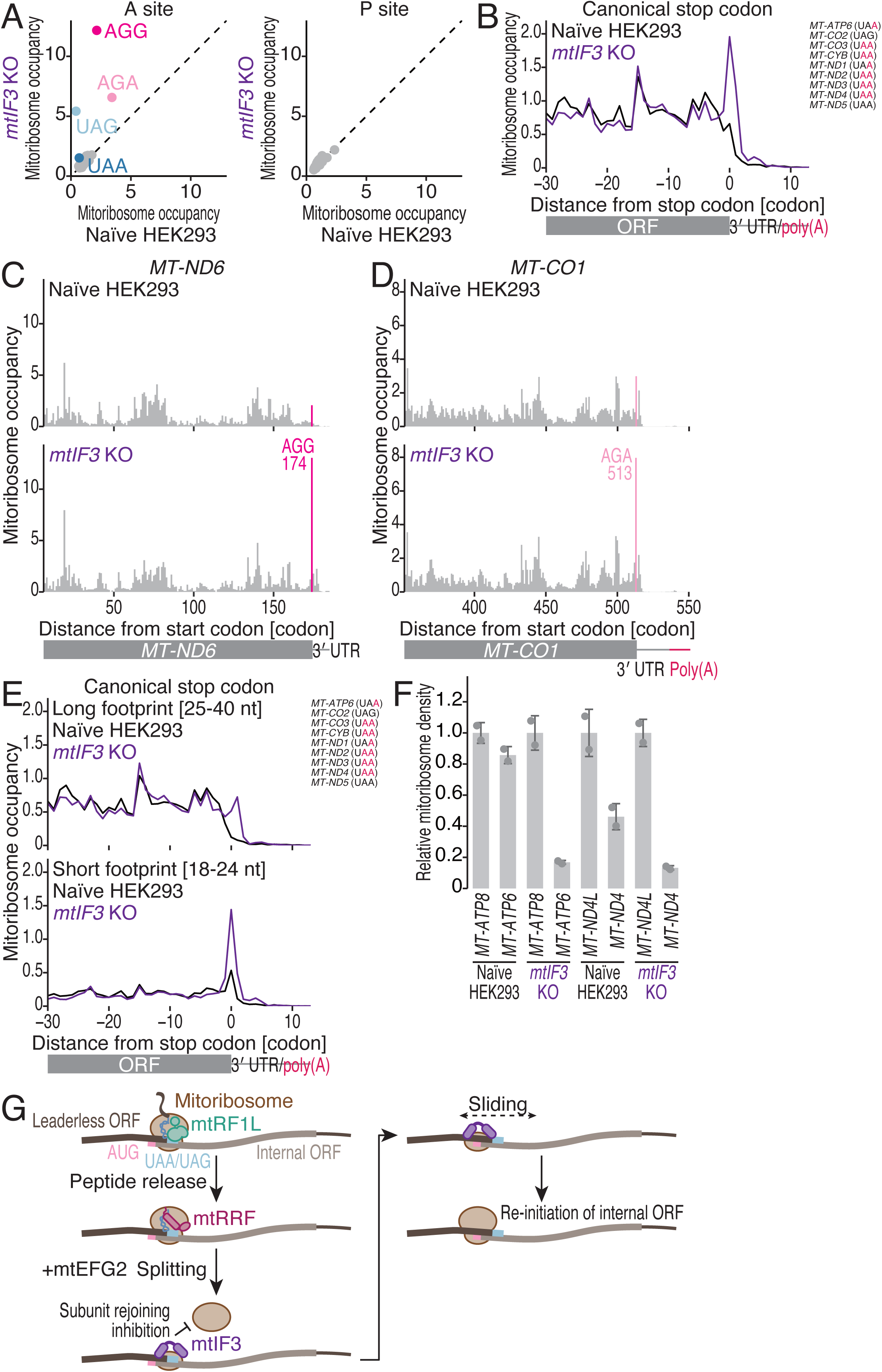
mtIF3 is required for mitoribosome recycling and internal ORF translation. (A) Comparison of mitoribosome occupancy at the A-site (left) and P-site (right) codons in the indicated cells. (B) Metagene plots of mitoribosome footprints around canonical stop codons in the indicated cells. The A-site position of each read is displayed. The ORFs and stop codon sequences that were used in the analysis are shown. The *MT-ATP8* and *MT-ND4L* ORFs were excluded from the analysis since the 3′ ends of these ORFs overlap with the downstream *MT-ATP6* and *MT-ND4* ORFs in the polycistronic mRNAs. (C and D) Distribution of mitoribosome footprints in naïve and *mtIF3* KO HeLa cells along the indicated transcripts. The A-site position of each read is displayed. The noncanonical stop codons are highlighted. (E) The same as B but with long footprints (25-40 nt, top) and short footprints (18-24 nt, bottom) separately plotted. (F) Relative mitoribosome density on the ORFs in the polycistronic mRNAs in the indicated cells. The data are normalized to the upstream ORFs (*MT-ATP8* and *MT-ND4L*) in the polycistronic mRNAs. The means (bars), s.d.s (errors), and individual replicates (n = 2, points) are shown. (G) Model of internal ORF translation in polycistronic mRNAs. See also Figure S7.

Together, these data support a model in which the reuse of mitoribosomes after termination of the translation of an upstream leaderless ORF facilitates the translation of the downstream internal ORF (Figure 5G). The requirement of mtRRF and mtIF3 suggested that after 39S dissociates, the mRNA-associated 28S subunit may reinitiate protein synthesis from internal ORFs (Figure 5G). Given that 40S cytosolic ribosomes may slide along mRNAs in both the 5′ and 3′ directions after termination and 60S dissociation ^54^, 28S mitoribosomes may follow the same movements, leading to recognition of AUG codons at the 5′ side from stop codons (Figure 5G).

### ICT1, mtRF-R, mtRES1, and mtRRF rescue mitoribosomes at start codons

We then focused on the class 2 release factors ICT1 and mtRF-R. Although structural and biochemical analyses have suggested that these proteins rescue stalled mitoribosomes ^15,21,22^, their endogenous substrates remain poorly understood. To address this issue, we generated KO HeLa cells in which *ICT1* and *mtRF-R* were knocked out (Figures S8A-D and S9A-D), and these cells may exhibit growth defects in a strain-dependent manner (Figures S8E and S9E) and reduced global mitochondrial translation (Figure S9F-G). Similar defects were observed in siRNA-mediated knockdown inHeLa cells ^24^ and patient-derived fibroblasts with a point mutation ^22^. Since mtRES1, which is an RNA-binding protein, forms a complex with mtRF-R to rescue stalled mitoribosomes ^21^, we also used *mtRES1* KO HAP1 cells, which are commercially available (Horizon Discovery) (Figure S10A); these cells exhibited mild cell growth inhibition (Figure S10B) and significant defects in mitochondrial translation (Figure S10C-D), as reported in HEK293 cells ^55^.

MitoIP-Thor-Ribo-Seq of these cell lines (Figures S8F-G, S9H-I, and S10E-F) revealed that the most striking molecular phenotype was the mitoribosome stalling at start codons (Figure 6A-C). The depletion of all the rescue factors caused the same pausing outcomes. The *MT-CO1* ORF is a remarkable example (Figures S8H, S9J, and S10G). Therefore, the distribution of the mitoribosomes was biased toward the 5′ end of the ORFs (Figures S8G, S9I, and S10F). In addition to leaderless ORFs, the start codons of internal ORFs exhibited mitoribosome accumulation, as illustrated by the results of *MT-ATP6* (Figure 6D-F). Focusing on this start codon, we observed a relative increase in the generation of short footprints, suggesting the accumulation of A site-free mitoribosomes (Figure 6D-F). Notably, the start codons of leaderless ORFs could not be used for this analysis because of the absence of the 5′ UTRs covered by mitoribosomes ^33,51^.

**Figure 6.**
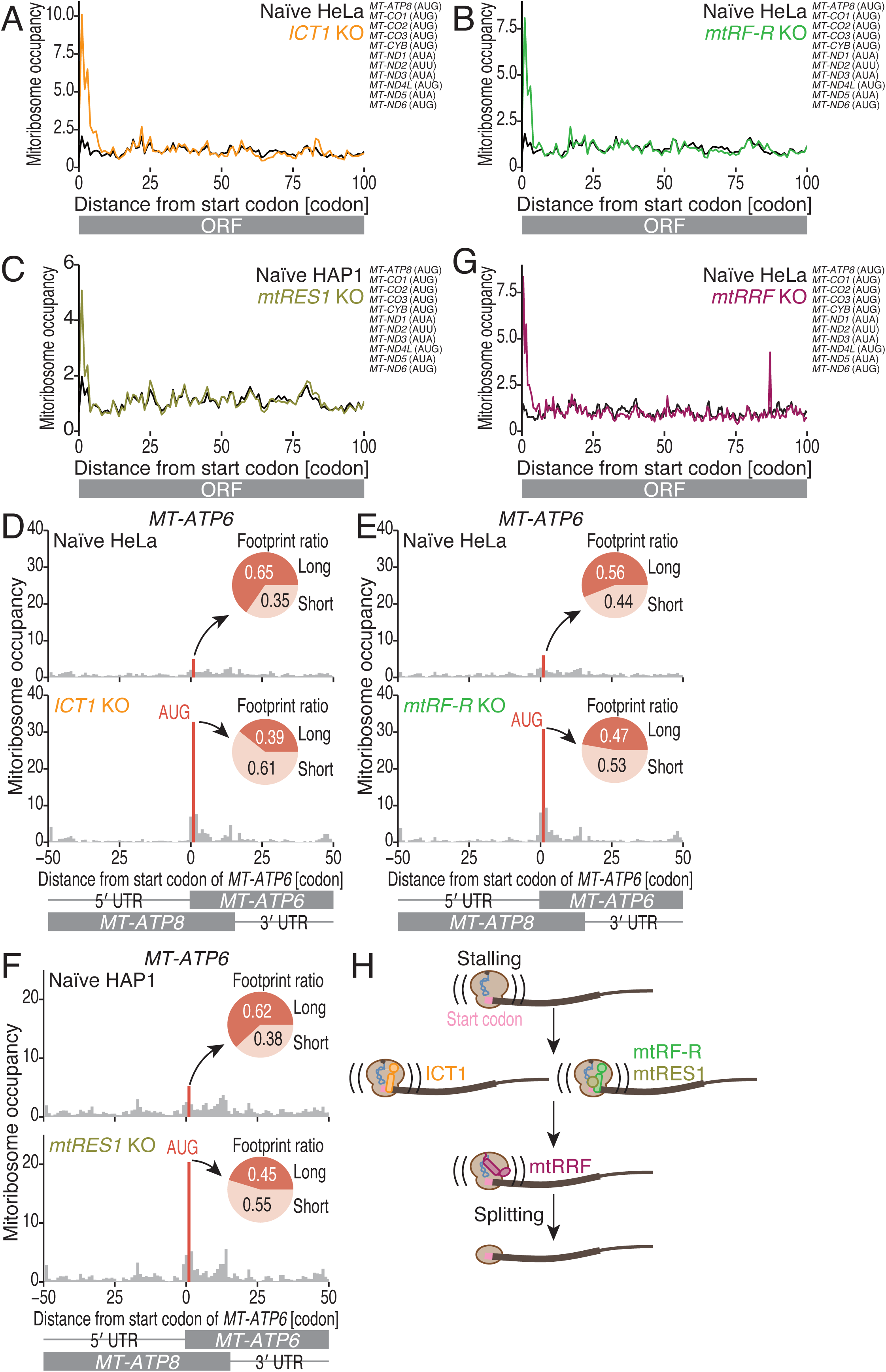
The start codon is a hot spot for quality control. (A-C and G) Metagene plots of mitoribosome footprints (the A site position) in naïve and *ICT1* KO HeLa cells (A), naïve and *mtRF-R* KO HeLa cells (B), naïve and *mtRES1* KO HAP1 cells (C), and naïve and *mtRRF* KO HeLa cells (G). (D-F) Distribution of mitoribosome footprints in naïve and *ICT1* KO HeLa cells (D), naïve and *mtRF-R* KO HeLa cells (E), and naïve and *mtRES1* KO HAP1 cells (F) along the indicated transcripts. The A-site position of each read is displayed. The inset pie chart represents the fractions of long footprints (25-40 nt) and short footprints (18-24 nt) at the start codon of *MT-ATP6*. (H) Model of quality control at the start codons. See also Figures S8-10.

Interestingly, a similar phenotype of mitoribosome traversal at start codons was observed in *mtRRF* KO HeLa cells (Figures 6G and S6P-Q). Although the cause of mitoribosome stalling remains unclear, our results suggested that mitoribosomes at the start codons are rescued by ICT1 and mtRF-R/mtRES1 and then recycled by mtRRF (Figure 6H).

Although the ICT1-mediated recognition of noncanonical stop codons has been a matter of debate ^19,20,23–25^, we did not observe mitoribosomes stalling at AGA/AGG codons in *ICT1* KO HeLa cells (Figure S8I). However, in *mtRF-R* KO HeLa (Figure S9K-L) and *mtRES*1 KO HAP1 cells (Figure S10H-I), we observed increased mitoribosome occupancy at the AGA codon of *MT-CO1*. The accumulation of mitoribosomes at the AGA codon was not as strong as observed under conditions of mtRF1 depletion, since we did not observe mitoribosome collision by MitoIP-Thor-Disome-Seq (Figures S8L-M, S9N-O, and S10K-L). Additionally, the mitoribosome footprints in the 3′ UTR downstream of the AGA codon were observed under conditions of mtRF-R (Figure S9L-M), mtRES1 (Figure S10I-J), and ICT1 (Figure S8J-K) depletion. The effects on mitoribosomes sliding or reading through in the 3′ UTR/poly(A) tail downstream of the canonical stop codons were also observed in *ICT1* and *mtRF-R* KO cells (Figures S8N-O, S9P-Q, and S10M). Thus, mitochondrial rescue factors may contribute to mitoribosome recycling and/or the suppression of stop codon readthrough in the 3′ UTR/poly(A) tail.

### Glutamine methylation of mitochondrial release factors supports efficient translation completion

Next, we investigated mechanisms by which termination reactions are regulated. Gln methylation of release factors at the catalytic GGQ motif is known to facilitate translation termination in bacteria (RF1 and RF2) ^56–59^ and in the cytosol of eukaryotes (eRF1) ^60–63^. The HEMK1-mediated methylation of Gln residues was observed in mtRF1, mtRF1L, ICT1, and mtRF-R ^21,64,65^. However, the impact of this methylation is still unclear.

Thus, MitoIP-Thor-Ribo-Seq was performed with *HEMK1* KO HeLa cells ^65^ to examine the role of GGQ methylation in mitochondrial translation (Figure S11A-B). We observed slight but detectable mitoribosome stalling on both canonical and noncanonical stop codons in response to HEMK1 depletion (Figures 7A-C and S11C-D). The re-expression of HEMK1 ^65^ counteracted this effect on termination (Figure 7A-C). In terms of read accumulation on the stop codons and 3′ UTR/poly(A) tail, the impact of HEMK1 depletion was milder than that of mtRF1L or mtRF1 depletion (Figures 1, 2, S1, and S2). Thus, we concluded that GGQ methylation is not a prerequisite for but rather enhances termination reactions by mtRF1L and mtRF1.

**Figure 7.**
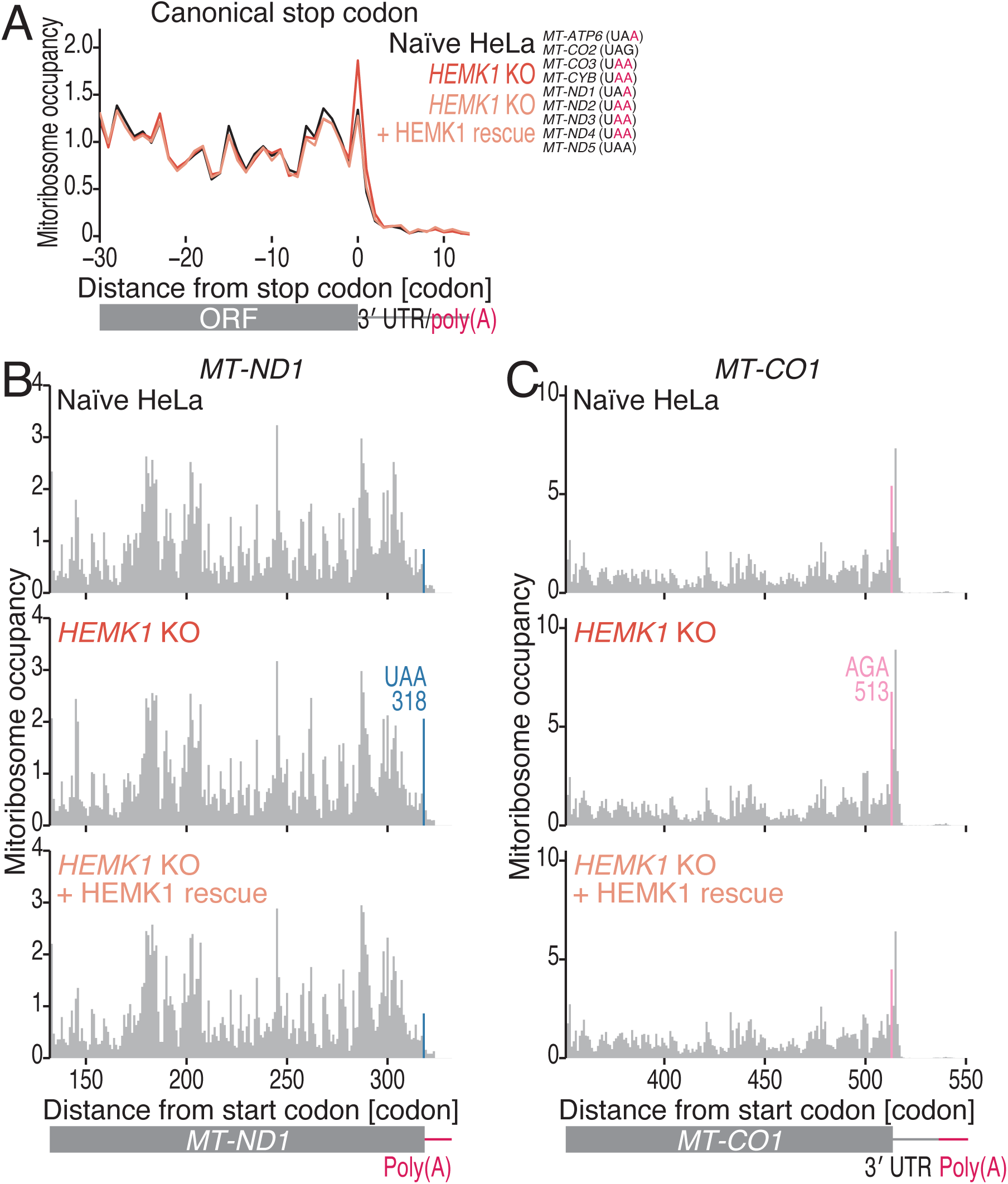
HEMK1-mediated GGQ methylation promotes translation termination. (A) Metagene plots of mitoribosome footprints around canonical stop codons in the indicated cells. The A-site position of each read is displayed. The ORFs and stop codon sequences that were used in the analysis are shown. The *MT-ATP8* and *MT-ND4L* ORFs were excluded from the analysis since the 3′ ends of these ORFs overlap with the downstream *MT-ATP6* and *MT-ND4* ORFs in the polycistronic mRNAs. (B and C) Distribution of mitoribosome footprints in naïve, *HEMK1*-KO, and *HEMK1* KO + HEMK1 rescue HeLa cells along the indicated transcripts. The A-site position of each read is displayed. The canonical and noncanonical stop codons are highlighted. See also Figure S11.

## Discussion

The high sensitivity of MitoIP-Thor-Ribo/Disome-Seq provides valuable insights into ongoing discussions about translation termination, mitoribosome recycling, reinitiation, and quality control (Figures 2G, 3G, 5G, and 6H). The utility of this techniques is exemplified by its ability to reveal the impact of HEMK1 on mitochondrial translation termination (Figure 7), which could not be assessed by previously available techniques ^65^. Our data are a useful resource and provide a global overview of mitochondrial protein synthesis.

Our data provide new information about the specificity of release factors for different kinds of stop codons (Figure 2G). The specificity of mtRF1L for canonical stop codons has been assessed with an *in vitro* translation system based on bacterial ribosomes ^13,14^ and metabolic labeling of newly synthesized proteins in cells ^16^. However, owing to the chimeric biochemical setup and radiolabeled detection, these approaches may not be able to detect the role of mtRF1L in the termination of translation at noncanonical stop codons. Our MitoIP-Thor-Ribo/Disome-Seq approach overcame these limitations and revealed the quantitative contribution of mtRF1L across stop codon sequences.

The role of mtIF3 in mitochondrial translation is still under debate. Considering its similarity to its bacterial homolog, mtIF3 was suggested to bind to 28S and then recruit mtIF2, fMet-tRNA^Met^, and leaderless mRNAs, ensuring correct tRNA usage ^66,67^. Alternatively, mtIF3 may dissociate from 28S before fMet-tRNA^Met^ and leaderless mRNA association, since 28S-bound mtIF3 may not be structurally compatible with fMet-tRNA^Met^ ^9,68^. Thus, direct binding of preassembled 55S to the start codon at the 5′ end of leaderless mRNA has been proposed ^53^. Importantly, mtIF3 may function differently in the translation of internal ORFs in polycistronic mRNAs; the loss of mtIF3 impacts the translation of *MT-ATP6* and *MT-ND4* more severely than the loss of other proteins ^33,52,53,66^. Our model of the recycling-coupled translation of internal ORFs (Figure 5G) is well aligned with the function of mtIF3 in mitoribosome recycling and the necessity of the translation of upstream ORFs for the translation of downstream internal ORFs ^32^.

Our results suggested a 28S sliding model for internal ORF translation (Figure 5G). Notably, the same mechanism should apply to translation from the cryptic initiation sites that were identified in a previous study ^33^. Importantly, whereas eukaryotic initiation factor 4F (eIF4F) — a complex of eIF4A, eIF4E, and eIF4G — provides 3′ directionality of 40S sliding ^54^, mitochondrial translation does not have such a factor to restrict 28S movement. Thus, 28S may have a better chance of recognizing start codons upstream of stop codons.

ICT1 has been suggested to recognize the mitoribosomes that stall at the end of a truncated mRNA with an empty A site ^15,19,20^. Recent structural analysis suggested that ArfB, which is a bacterial homolog of ICT1, may displace mRNA at the A site ^69,70^. Thus, mitoribosomes that stall at a start codon may serve as substrates of ICT1 and mtRF-R.

## Limitations

We observed footprints from mitoribosomes reaching the 3′ UTR/poly(A) regions when mtRF1, mtRF1L, mtRRF, ICT1, and mtRF-R were knocked out (Figures 1, 2, 4, 5, S1, S2, S5, S6, S8, and S9). Some fractions could represent unrecycled mitoribosomes, particularly in *mtRRF* KO cells (Figure 4). Even without the depletion of these genes, the AGA noncanonical stop codon in *MT-CO1* was originally weak and generated a fraction of unrecycled mitoribosomes and the subsequent collisions of mitoribosomes ^33^. The other mitoribosome fraction downstream of stop codons may have emerged due to stop codon readthrough. The situation is more straightforward in cells lacking mtRF1 and mtRF1L, considering the outcomes of cytosolic eukaryotic translation release factor 1 (eRF1) depletion and observations made in studies of release factor-deficient bacterial strains ^40,41^. This study is limited in that it explicitly investigated the states of mitoribosomes downstream of the stop codons.

Our data suggested substantial protein synthesis from out-of-frame sequences (Figures 3, S3, and S4). In addition to secondary RNA structure-associated frameshifts, another potential source could be cryptic translation initiation, as suggested by another study ^33^. Further work is needed to identify the source and cause of out-of-frame translation.

## Acknowledgments

We thank all the members of the Iwasaki laboratory and Dr. Asuteka Nagao for constructive discussion and technical help. S.I. was supported by the Ministry of Education, Culture, Sports, Science and Technology (MEXT) (JP20H05784 and JP24H02307), the Japan Society for the Promotion of Science (JSPS) (JP23H02415 and JP23H00095), the Japan Agency for Medical Research and Development (AMED) (JP20gm1410001), the Gushinkai Foundation, and RIKEN [Pioneering project “Biology of Intracellular Environments” and RIKEN TRIP initiative (TRIP-AGIS)]. T.W. was supported by The Graduate School of Frontier Sciences, The University of Tokyo (C2205 and C2306) and JSPS (JP23KJ0444 and JP24K23198). Y.S. was supported by a RIKEN intenal research fund. Computations were supported by the supercomputer HOKUSAI SailingShip in RIKEN. We are grateful to the Support Unit for Bio-Material Analysis, RIKEN CBS Research Resources Division, for Sanger sequencing and cell sorting. T.W. was a recipient of fellowships from JSPS (DC2) and JST SPRING (JPMJSP2108). Q.F. was an International Program Associate of RIKEN. A part of the DNA libraries was sequenced at the Advanced Genomics Center, National Institute of Genetics, Mishima, Shizuoka, Japan.

## Author contributions

Conceptualization, T.W., M.M., and S.I.;

Methodology, T.W., M.M., Q.F., and Y.I.; Formal analysis, T.W.;

Investigation, T.W., M.M., Q.F., and Y.I.;

Resources, Q.F. and Y.S.;

Writing – Original Draft, S.I.;

Writing – Review & Editing, T.W., M.M., Q.F., Y.I., Y.S., and S.I.;

Visualization, T.W., Y.I., and S.I.;

Supervision, Y.I., Y.S., and S.I.;

Project administration, S.I.;

Funding Acquisition, T.W., Y.S., and S.I.

## Materials and Methods

### Cell culture

Naïve HeLa cells (RIKEN BioResource Research Center), HEK293 cells [American Type Culture Collection (ATCC), CRL-1573], HAP1 cells (Horizon Discovery, C631), and C2C12 cells (ATCC, CRL-1772) were cultured at 37°C in 5% CO_2_ in the media described in Table S1. An e-Myco VALiD Mycoplasma PCR Detection Kit (iNtRON Biotechnology) was used to maintain *Mycoplasma*-free cultures.

### KO cell lines

*HEMK1* KO HeLa cells, *HEMK1* KO + HEMK1 rescue HeLa cells, and *mtIF3* KO HEK293 cells were previously established ^33,65^. *mtRES1* KO HAP1 cells were purchased from Dhamacon (Horizon Discovery, 5-bp deletion, HZGHC004965c010).

*mtRF1L*, *mtRF1*, *mtRRF*, *ICT1*, and *mtRF-R* KO HeLa cells and *mtRF1* KO C2C12 cells were established via CRISPR-Cas9–mediated genome editing according to a previously described protocol ^71^. Each pair of spacer sequences in the guide RNAs was designed to target the exons of human *mtRF1L*, human *mtRF1*, human *mtRRF*, human *ICT1*, human *mtRF-R*, or mouse *mtRF1* (human *mtRF1L*, 5′-GACAUGCAUCUGCCAGCAUU-3′ and 5′-CGGAUAGUUCAUCUUCCAAC-3′; human *mtRF1*, 5′-CCAUGCCUUCUGUUCUAGG-3′ and 5′-AGACUUGUCCACAUCCCCAC-3′; human *mtRRF*, 5′-AGUACAAAGCUUGCAACUUC-3′ and 5′-ACGUCACAUGUCUCUACCUU-3′; human *ICT1*, 5′-AUAUCUUAUUGUCGGAGUAG-3′ and 5′-AUGUUAAACUUCAUAGAAUC-3′; human *mtRF-R*, 5′-CACGUUCCUUAUGAGCACCG-3′ and 5′-CCCCUCAGGCAUCGUUGUAA-3′; and mouse *mtRF1*, 5′-AAAGCAAUUCUAAAGAUAUA-3′ and 5′-AUUGUGUAAAAGUAGGUAAA-3′). These spacer sequences were subsequently cloned and inserted into the pL-CRISPR.EFS.GFP (a gift from Benjamin Ebert, Addgene plasmid #57818; http://n2t.net/addgene:57818; RRID: Addgene_57818) and pL-CRISPR.EFS.tRFP (a gift from Benjamin Ebert, Addgene plasmid #57819; http://n2t.net/addgene:57819; RRID: Addgene_57819) ^72^. Both plasmids expressing guide RNAs were subsequently transfected into naïve HeLa cells or naïve C2C12 cells with Lipofectamine 3000 (Thermo Fisher Scientific) according to the manufacturer’s instructions. Forty-eight hours after transfection, flow cytometry was performed with a FACSAria II Special Order system (BD) to sort the GFP and RFP double-positive cell populations. After the recovery culture, the clones were isolated by limited dilution cloning in 96-well plates. Gene knockout was confirmed by Sanger sequencing of genomic DNA and Western blotting.

### Lysate preparation for MitoIP-Thor-Ribo-Seq and MitoIP-Thor-Disome-Seq

Lysate preparation was performed according to a previously described protocol ^33^. Cells were washed with ice-cold PBS and lysed with hypotonic buffer (10 mM HEPES-KOH pH 7.5, 10 mM KCl, 1.5 mM MgCl_2_, 1 mM DTT, 100 µg/ml cycloheximide, and 100 µg/ml chloramphenicol). Then, mitochondrial purification was performed by immunoprecipitation with anti-TOM22 antibody-conjugated beads from a Mitochondria Isolation Kit, Human (Miltenyi Biotec) or a Mitochondria Isolation Kit, mouse tissue (Miltenyi Biotec). After the washing and elution steps, the mitochondrial fraction was pelleted by centrifugation at 7,000 × g for 10 min at 4°C and resuspended in modified lysis buffer (20 mM Tris-HCl pH 7.5, 150 mM NaCl, 15 mM MgCl_2_, 1 mM DTT, 1% Triton X-100, 100 µg/ml cycloheximide, 100 µg/ml chloramphenicol, and 2.5 U/ml TURBO DNase). The lysate was clarified by centrifugation at 20,000 × *g* for 10 min at 4°C.

### Library preparation for MitoIP-Thor-Ribo-Seq and MitoIP-Thor-Disome-Seq

The library was prepared as previously reported ^33^. The mitochondria-enriched lysate was subjected to RNase digestion, sucrose cushion ultracentrifugation, and Ribo-FilterOut ^73^. Then, mitoribosome footprints ranging from 17 to 50 nt in length for the mito-monosome footprints and from 50 to 80 nt in length for the mito-disome footprints were gel-excised.

After linker ligation, rRNA depletion was performed with a Ribo-Zero Gold rRNA Removal Kit (Human/Mouse/Rat) (Illumina, accompanied by a TruSeq Stranded Total RNA Kit), a Human Ribo-Seq riboPOOL kit (siTOOLs Biotech), or a Mouse/Rat Ribo-Seq riboPOOL kit (siTOOLs Biotech). An oligonucleotide was hybridized to the T7 promoter region of the linker to produce the double-stranded DNA T7 promoter. Antisense RNAs were transcribed with a T7-Scribe Standard RNA IVT Kit (CELLSCRIPT). After ligation of the second linker, the cDNA was synthesized by reverse transcription and amplified by PCR. DNA libraries were sequenced on a HiSeq X or a NovaSeq X Plus platform in 150-bp paired-end mode or on a NovaSeq 6000 platform in 100-bp single-end mode.

### Data analysis

For pair-end sequencing, base correction was performed using fastp (version 0.21.0) ^74^. Read 1 was processed for downstream analysis. After read quality filtering and adapter sequence removal by fastp, all the reads were aligned to noncoding RNAs (rRNAs, tRNAs, mt-rRNAs, mt-tRNAs, snRNAs, snoRNAs, and miRNAs). The remaining reads were subsequently mapped to the human or mouse nuclear genomes (human, hg38; mouse, mm10) and custom-made mitochondrial transcript sequences using STAR (version 2.7.0a) ^75^. Read suppression by a unique molecular identifier (UMI) was performed with UMItools (version 1.1.2) ^76^. Mitoribosomal A-site offsets for each footprint length were empirically estimated as previously described (Table S1) ^33^.

To calculate mitoribosome occupancy, the reads of each codon were normalized by the average reads per codon of the ORF. The first 5 codons of each transcript and overlapping ORF regions in the bicistronic transcripts were omitted from the calculation (except where indicated in the figure panels and legends).

The polarity score was defined as previously reported ^77^.

### On-gel mito-FUNCAT

On-gel mito-FUNCAT was performed according to a previously described protocol ^33,71,78^. Cells were cultured in 6-well plates in methionine-free medium supplemented with 50 µM HPG (Jena Bioscience) and 100 µg/ml anisomycin (Alomone Labs) for 3 h before the cells were harvested. After the cells were lysed, the nascent peptides were labeled with 50 μM IRdye800CW Azide (LI-COR Biosciences) using a Click-it Cell Reaction Buffer Kit (Thermo Fisher Scientific) according to the manufacturer’s instructions. The reactants were separated on SDS–PAGE gels, and the signals were detected with an Odyssey CLx (LI-COR Biosciences) with an IR 800-nm channel. The signals were standardized against Coomassie Brilliant Blue (CBB) staining (GelCode Blue Safe Protein Stain, Thermo Fisher Scientific), which was detected by an IR 700-nm channel in an Odyssey CLx. The images were quantified with Image Studio (LI-COR Biosciences, version 5.2).

### Western blotting

Anti-β-actin [MEDICAL & BIOLOGICAL LABORATORIES (MBL), M177-3, 1:1000], anti-α-Tubulin (CST, 2144, 1:1000), anti-TOM20 [Cell Signaling Technology (CST), 42406, 1:1000], anti-mtRF1L (Proteintech, 16694-1-AP, 1:1000), anti-mtRF1 (Thermo Fisher Scientific, PA5-97988, 1:1000; Atlas Antibodies, HPA043316, 1:1000), anti-mtRRF (Proteintech, 12357-2-AP, 1:1000), anti-MT-CO1 (Abcam, ab203912, 1:200), anti-MT-CO2 (Proteintech, 55070-1-AP, 1:1000), anti-MT-ATP8 (Proteintech, 26723-1-AP, 1:1000), anti-MT-ATP6 (CST, 70262, 1:200), anti-MT-CYB (Proteintech, 55090-1-AP, 1:1000), anti-MT-ND4L (Thermo Fisher Scientific, PA5-68242, 1:200), anti-MT-ND4 (Merck Millipore, MABS1994, 1:200), anti-MRPS22/mS22 (Thermo Fisher Scientific, PA5-52249, 1:1000), anti-MRPL45/ml45 (Thermo Fisher Scientific, PA5-54784,1:1000), anti-ICT1 (Proteintech, 10403-1-AP, 1:1000), anti-mtRF-R (Proteintech, 24646-1-AP, 1:1000), and anti-mtRES1 (Abcam, ab151066, 1:1000) antibodies were used as primary antibodies. IRDye800CW-conjugated anti-rabbit IgG (LI-COR Biosciences, 926-32211, 1:10000) or IRDye800CW-conjugated anti-mouse IgG (LI-COR Biosciences, 926-32210, 1:10000) was used as a secondary antibody. Images were acquired with an Odyssey CLx (LI-COR Biosciences), and protein levels were quantified with Image Studio (LI-COR Biosciences, version 5.2).

### Mitochondrial purification

Mitochondrial purification was performed as described above (see the “*MitoIP-Thor-Ribo-Seq and MitoIP-Thor-Disome-Seq*” section). Cycloheximide and chloramphenicol were excluded from all the solutions. The supernatant was subjected to Western blotting.

### Cell viability assay

Cells were cultured in 96-well plates with or without supplements [1 × Na-pyruvate (nacalai tesque) and 0.05 g/l uridine]. Then, the cells were incubated with RealTime-Glo MT Cell Viability Assay reagent (Promega). Luminescence was monitored by the GloMax Navigator System (Promega).

### Modeling structures

The modeled structures of the interactions between the AGA and AGG stop codons and mtRF1L in the mitoribosome were generated manually with Coot ^79^ on the basis of the reported cryo-EM structure of the mitoribosomal termination complex of mtRF1L with the UAA codon (PDB ID: 7NQH) ^15^. The UAA nucleotides were replaced by AGA and AGG, and the neighboring mtRF1 residues were manually fitted to these nucleotides, followed by local regularization. ChimeraX ^80^ was used to prepare the figures.

## Data and code availability

The results of MitoIP-Thor-Ribo-Seq and MitoIP-Thor-Disome-Seq (GEO: GSE288420) that were obtained in this study were deposited in the National Center for Biotechnology Information (NCBI) database. The key custom scripts used in this study are available at Zenodo (https://zenodo.org, DOI: 10.1101/2023.07.19.549812).

---

**Figure S1.**
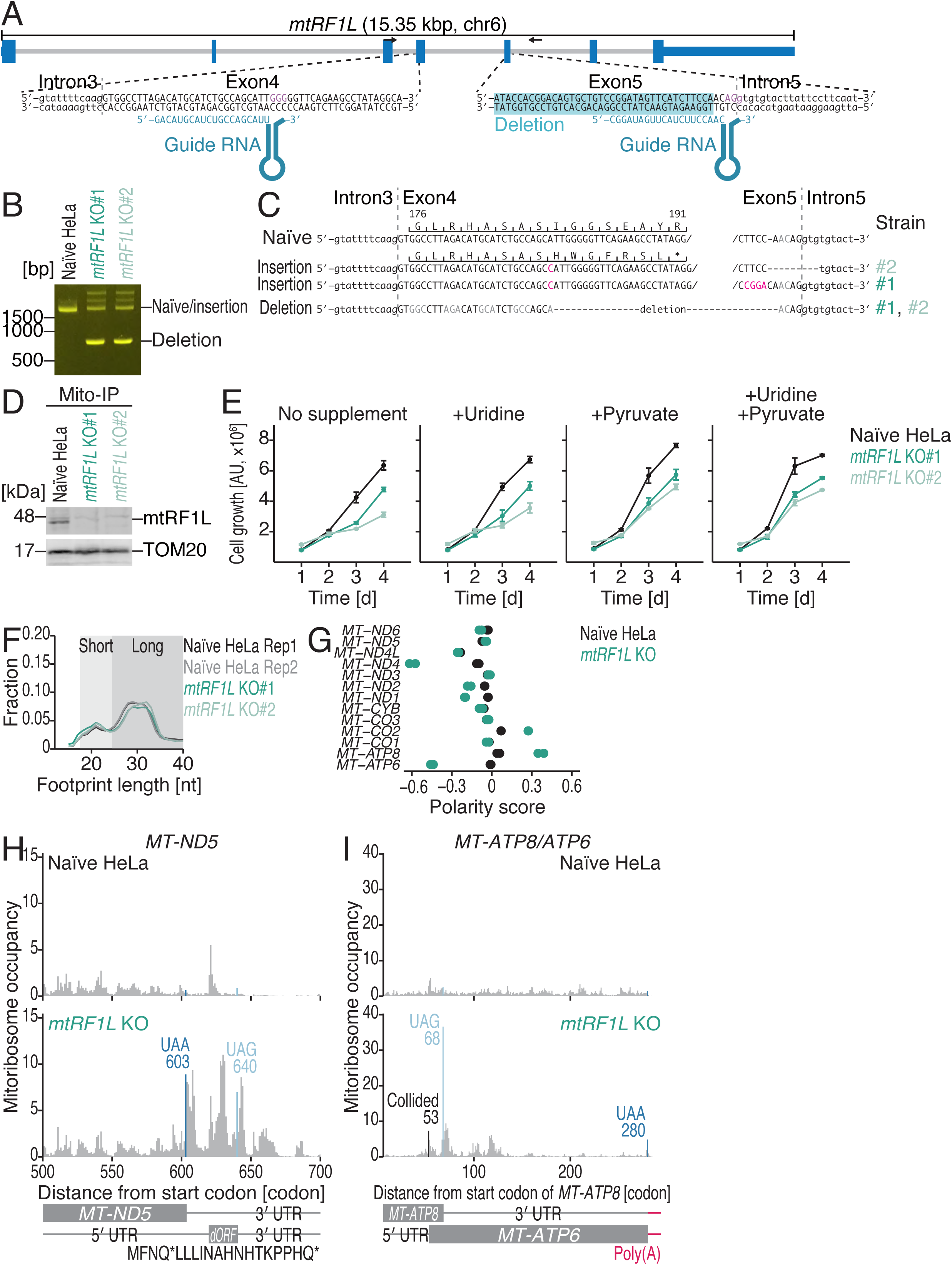
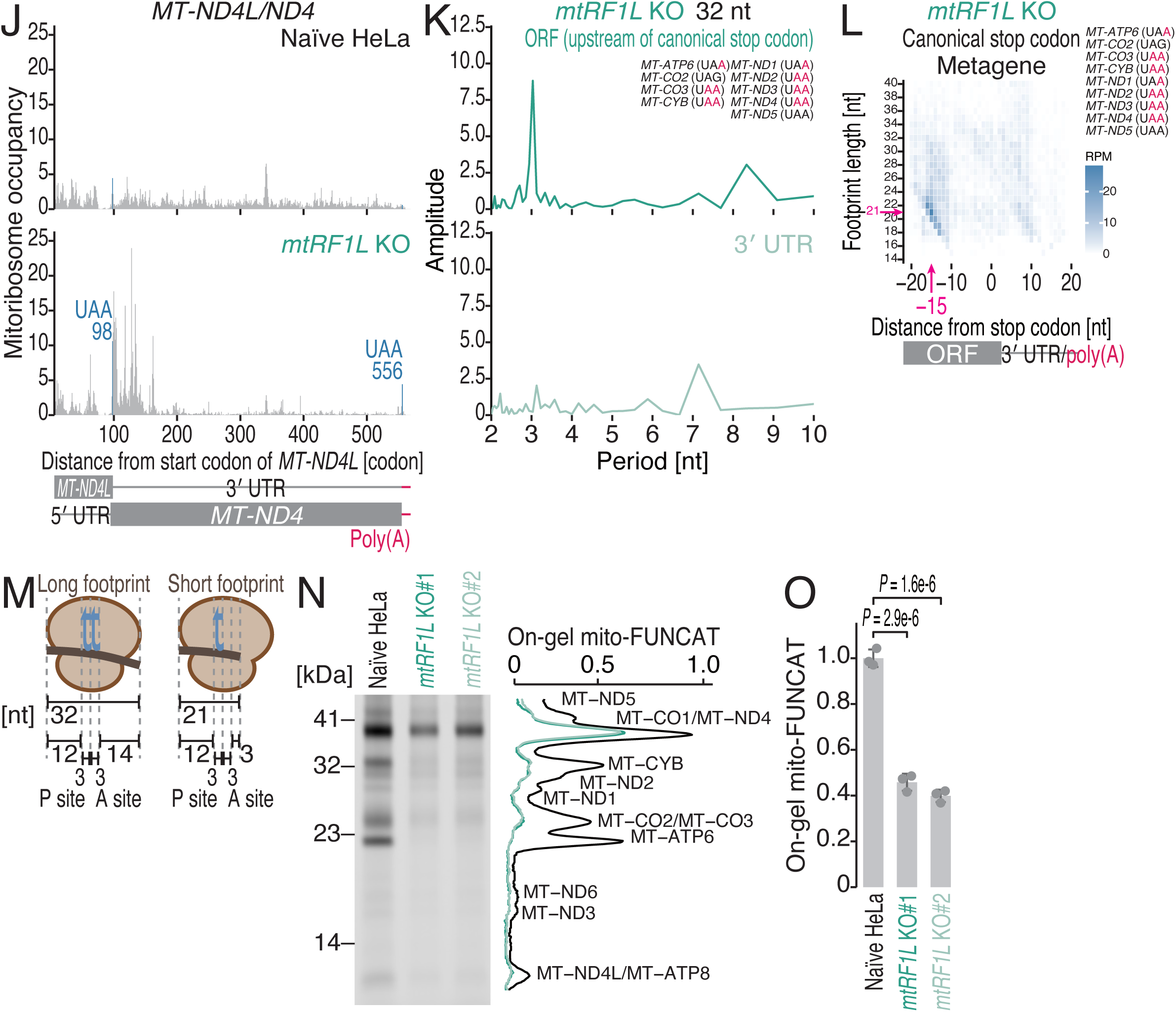
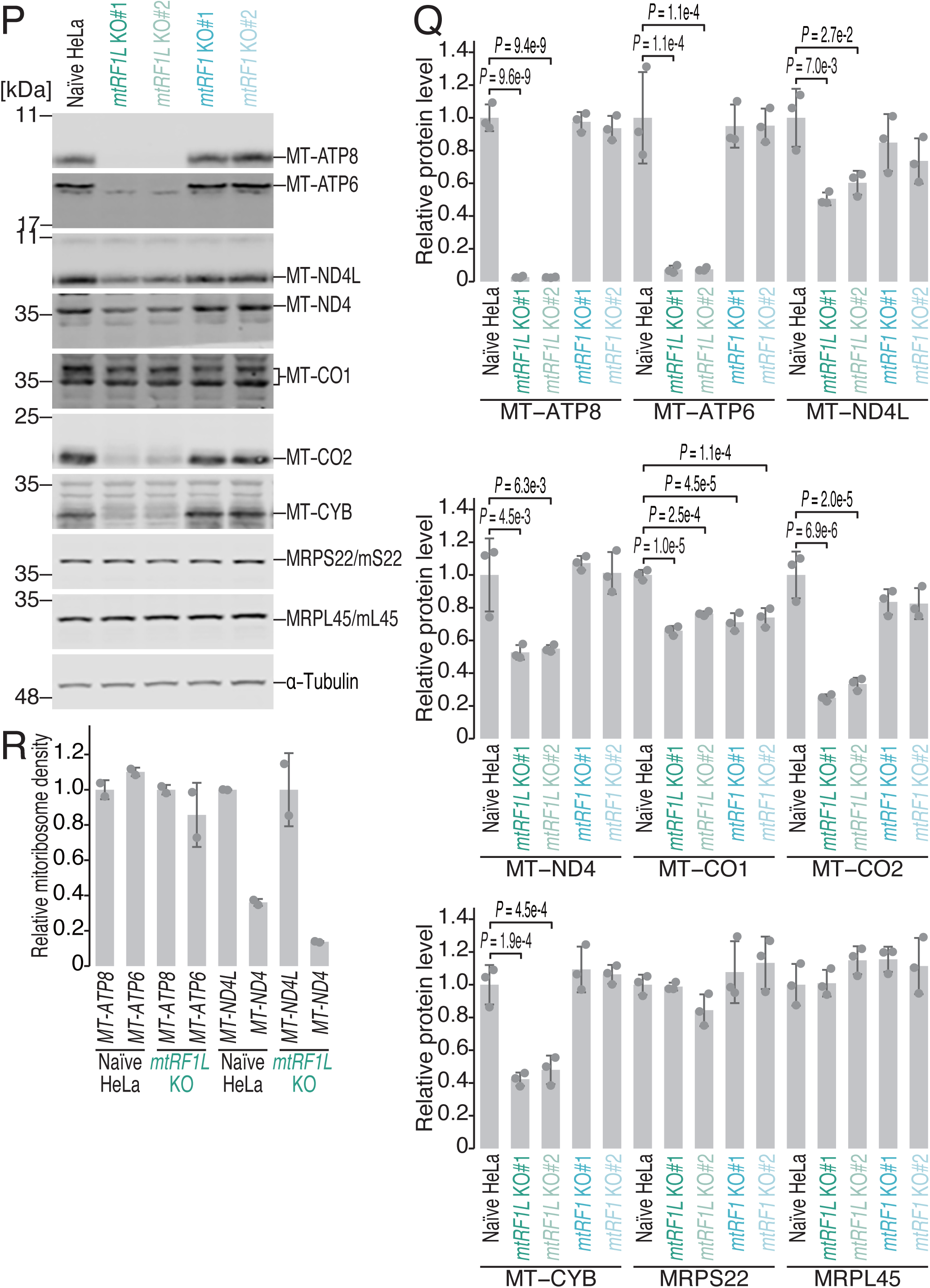
Characterization of MitoIP-Thor-Ribo-Seq data from *mtRF1L* KO HeLa cells, related to Figure 1. (A) Schematics showing the guide RNAs that were designed for CRISPR-Cas9-mediated gene knockout of human *mtRF1L* and the deleted region in the isolated cell lines. (B) Genomic PCR of the *mtRF1L* gene locus in the isolated cell lines. The primer positions are shown in A. (C) Schematic of the genomic alterations in the isolated cell lines. (D) The loss of the mtRF1L protein in the isolated cell lines was confirmed by Western blotting. The mitochondria-enriched fraction was subjected to immunoprecipitation with an anti-TOM22 antibody (MitoIP). TOM20 was used as a loading control. (E) Growth assay with naïve and *mtRF1L* KO HeLa cells grown under the indicated conditions. The means (points) and s.d.s (errors, n = 3) are shown. (F) The fraction of read lengths of mitoribosome footprints in the indicated cells. Rep, replicate. (G) Polarity scores of the indicated ORFs in the indicated cells. The data points from the two replicates are shown individually. (H-J) Distribution of mitoribosome footprints in naïve and *mtRF1L* KO HeLa cells along the indicated transcripts. The A-site position of each read is displayed. The canonical stop codon and the codon position indicating potential colliding mitoribosomes are highlighted. (K) Discrete Fourier transform of mitoribosome footprints mapped to the 100-nt region upstream and downstream of canonical stop codons. The ORFs and stop codon sequences that were used in the analysis are shown. (L) Plots of the 5′ ends of the mitoribosome footprint along the length around canonical stop codons (the first nucleotide of the stop codon was set to 0). The ORFs and stop codon sequences that were used in the analysis are shown. The color scale indicates read abundance. (M) Schematic representation of the relative positions of the P site and A site on the long (32 nt) and short (21 nt) mitoribosome footprints. (N) Gel image of on-gel mito-FUNCAT experiments performed with the indicated cell lines. Proteins newly synthesized by mitoribosomes are labeled with L-homopropargyl glycine (HPG) and conjugated with infrared 800 (IR800) dye (left). The signal intensities along the lanes are shown (right). (O) Quantification of the on-gel mito-FUNCAT data in N. The means (bars), s.d.s (errors), and individual replicates (n = 3, points) are shown. CBB staining of total proteins was used for data normalization. The p values were calculated with the Tukey-Kramer test (two-tailed). (P) Western blotting for the indicated proteins in the indicated cell lines. α-Tubulin was used as a loading control. (Q) Quantification of the on-gel mito-FUNCAT data in P. The means (bars), s.d.s (errors), and individual replicates (n = 3, points) are shown. α-Tubulin was used for data normalization. The p values were calculated with the Tukey‒Kramer test (two-tailed). (R) Relative mitoribosome density on the ORFs of the polycistronic mRNAs in the indicated cells. The data are normalized to the upstream ORFs (*MT-ATP8* and *MT-ND4L*) in the polycistronic mRNAs. The means (bars), s.d.s (errors), and individual replicates (n = 2, points) are shown.

**Figure S2.**
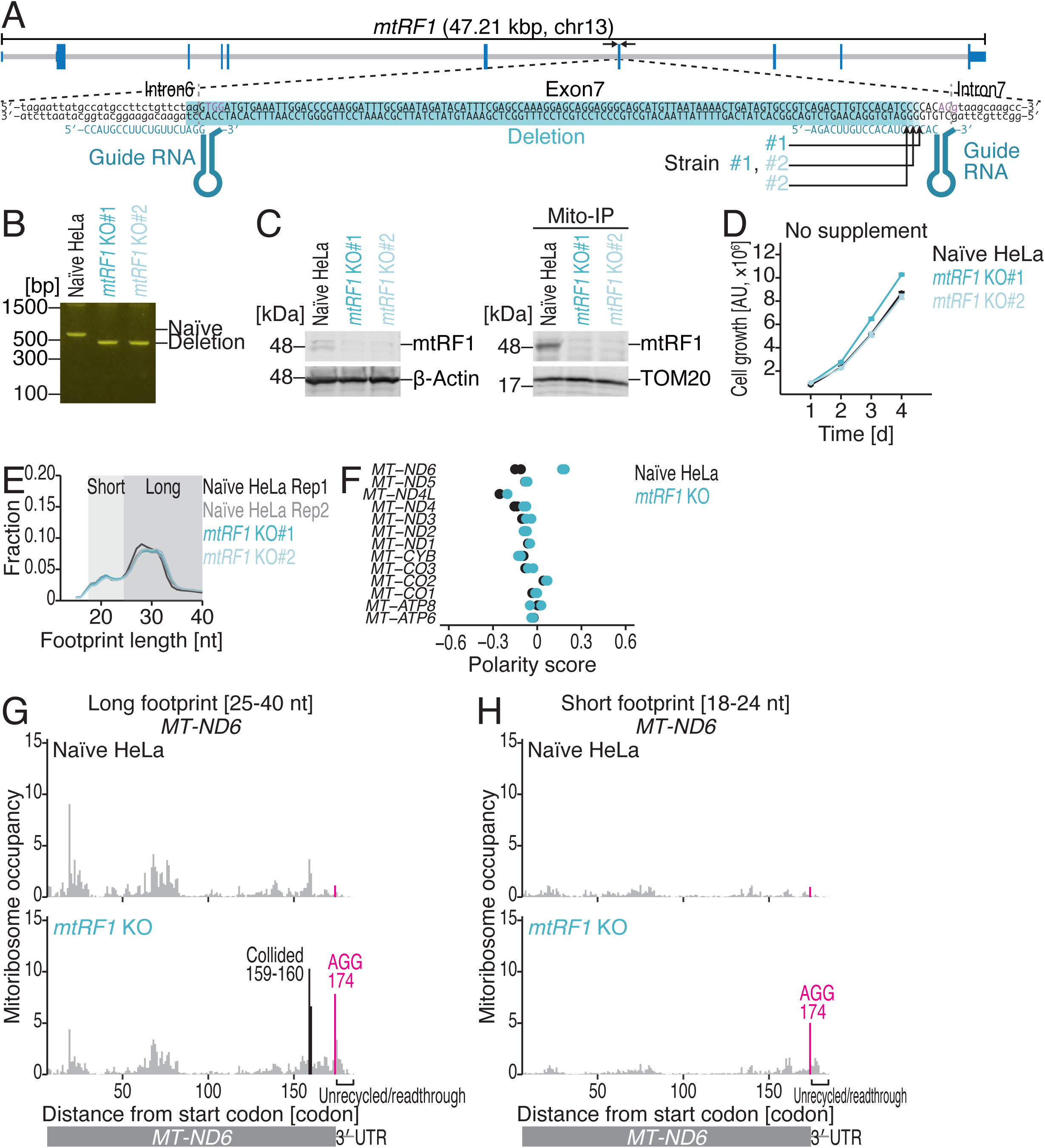
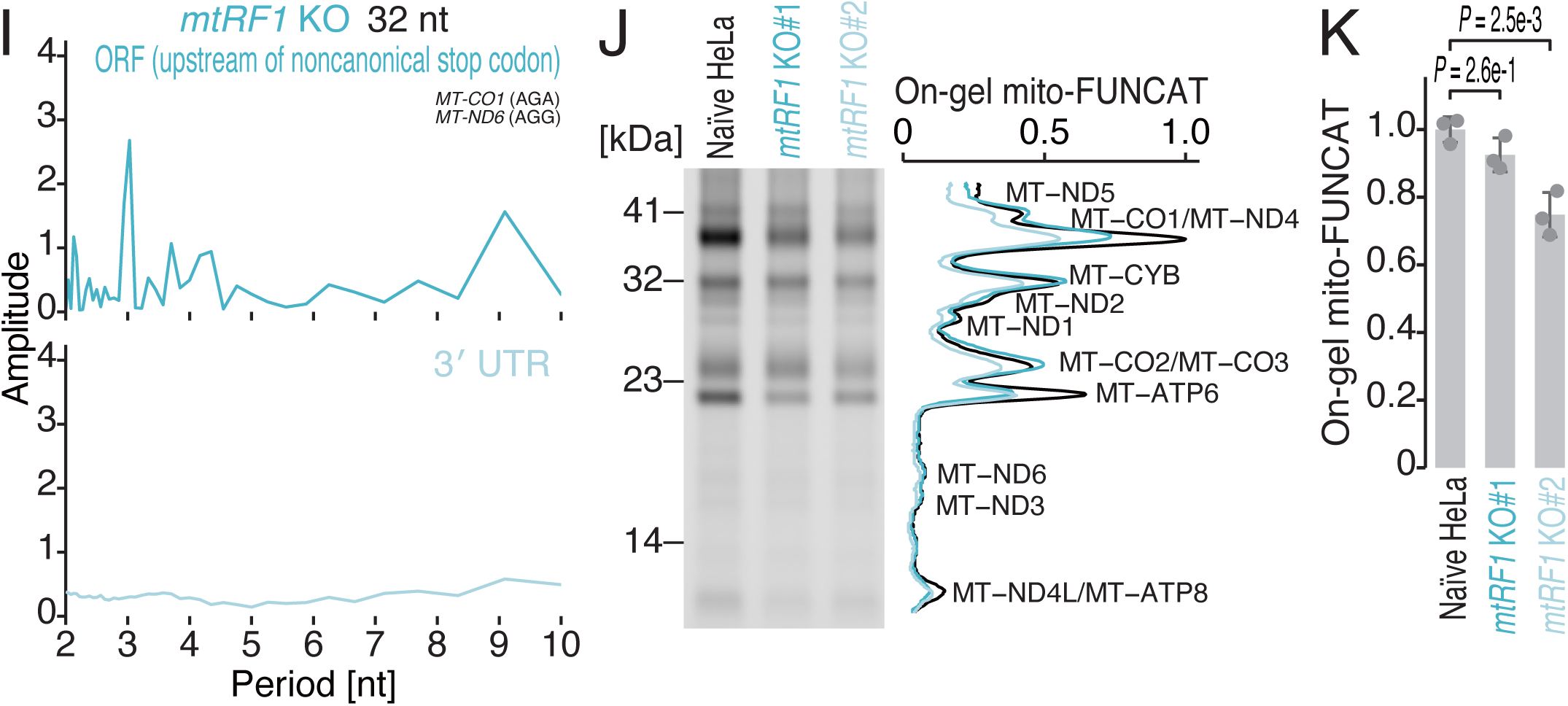
Characterization of MitoIP-Thor-Ribo-Seq data from *mtRF1* KO HeLa cells, related to Figure 2. (A) Schematics showing the guide RNAs that were designed for CRISPR-Cas9-mediated gene knockout of human *mtRF1* and the deleted region in the isolated cell lines. The genomic alterations in the isolated cell lines are shown. (B) Genomic PCR of the *mtRF1* gene locus in the isolated cell lines. The primer positions are shown in A. (C) The loss of the mtRF1 protein in the isolated cell lines was confirmed by Western blotting. Total lysates and mitochondria-enriched fractions obtained via immunoprecipitation with an anti-TOM22 antibody (MitoIP) were used. β-Actin and TOM20 were used as loading controls. (D) Growth assay with naïve and *mtRF1* KO HeLa cells grown under the indicated conditions. The means (points) and s.d.s (errors, n = 3) are shown. (E) The fraction of read length of mitoribosome footprints in the indicated cells. Rep, replicate. (F) Polarity scores of the indicated ORFs in the indicated cells. The data points from the two replicates are shown individually. (G and H) Distribution of mitoribosome footprints in naïve and *mtRF1* KO HeLa cells along the indicated transcripts. The A-site position of each read is displayed. The noncanonical stop codon and the codon position indicating potential colliding mitoribosomes are highlighted. (I) Discrete Fourier transform of mitoribosome footprints mapped to the 100-nt region upstream and downstream of noncanonical stop codons. The ORFs and stop codon sequences that were used in the analysis are shown. (J) Gel image of on-gel mito-FUNCAT experiments performed with the indicated cell lines. Proteins newly synthesized by mitoribosomes are labeled with L-homopropargyl glycine (HPG) and conjugated with infrared 800 (IR800) dye (left). The signal intensities along the lanes are shown (right). We note that the data for the naïve samples are the same as those in Figure S1N. (K) Quantification of the on-gel mito-FUNCAT data in J. The means (bars), s.d.s (errors), and individual replicates (n = 3, points) are shown. CBB staining of total proteins was used for data normalization. The p values were calculated with the Tukey‒Kramer test (two-tailed). We note that naïve samples are the same data used for Figure S1O.

**Figure S3.**
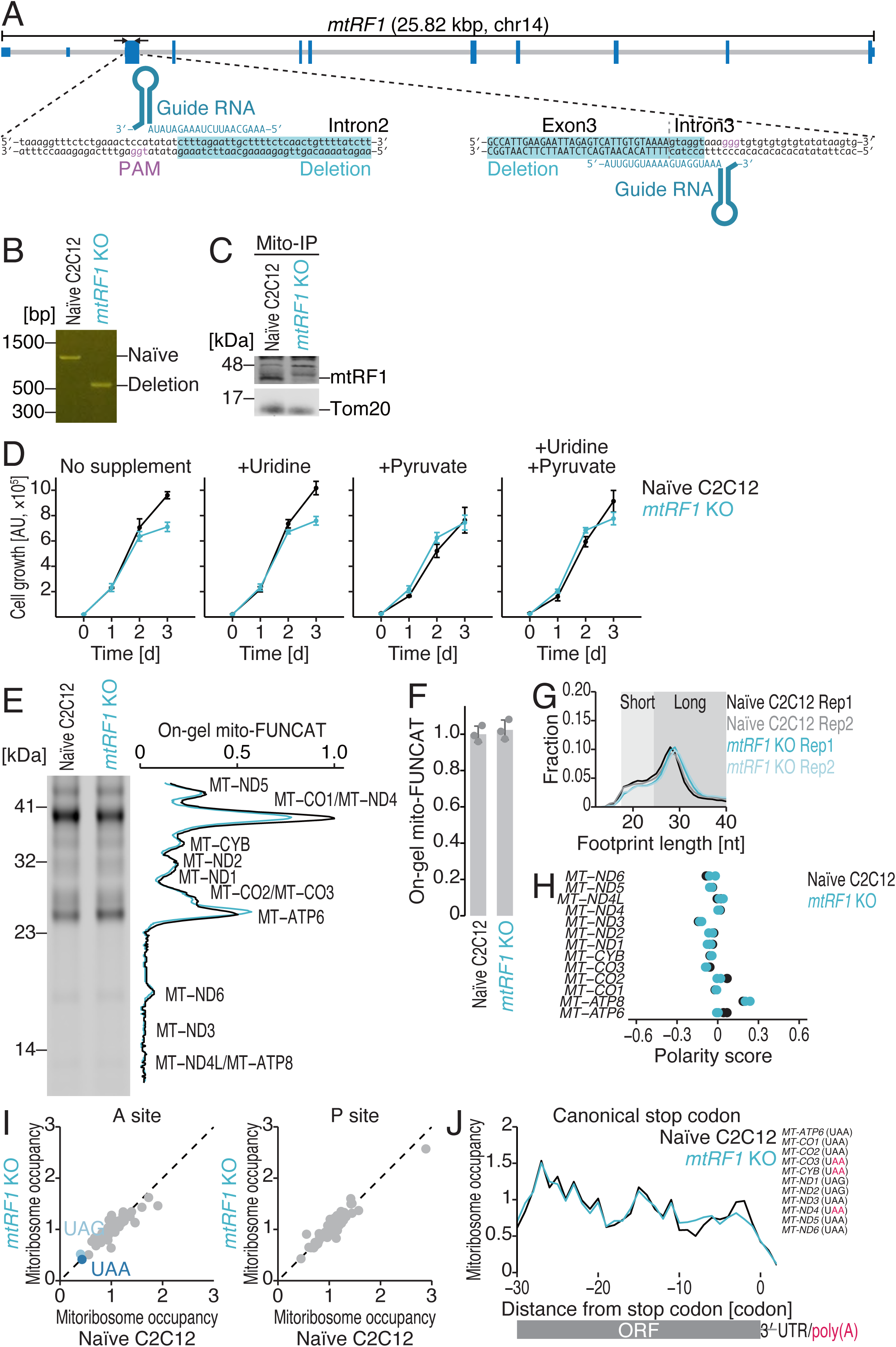
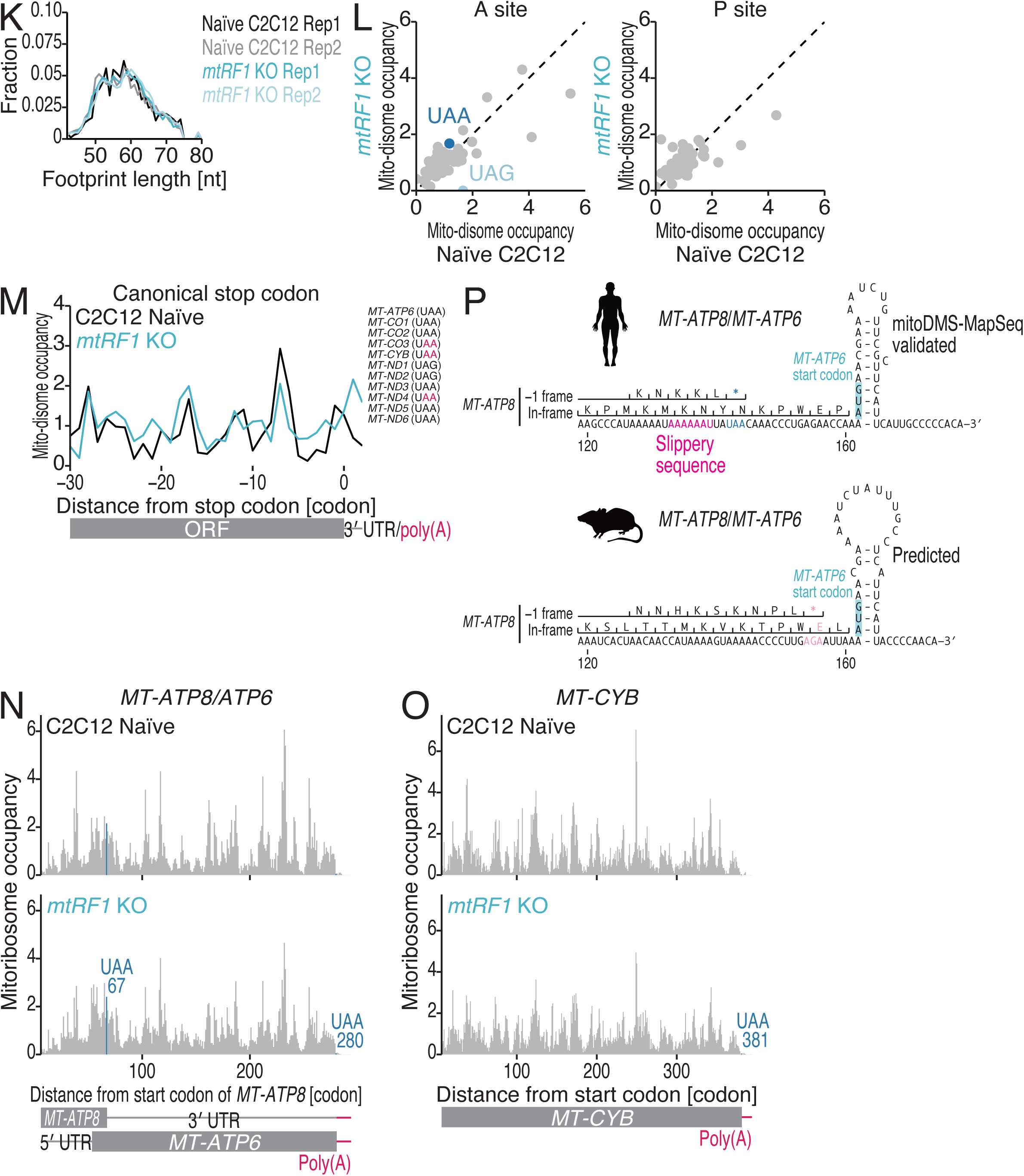
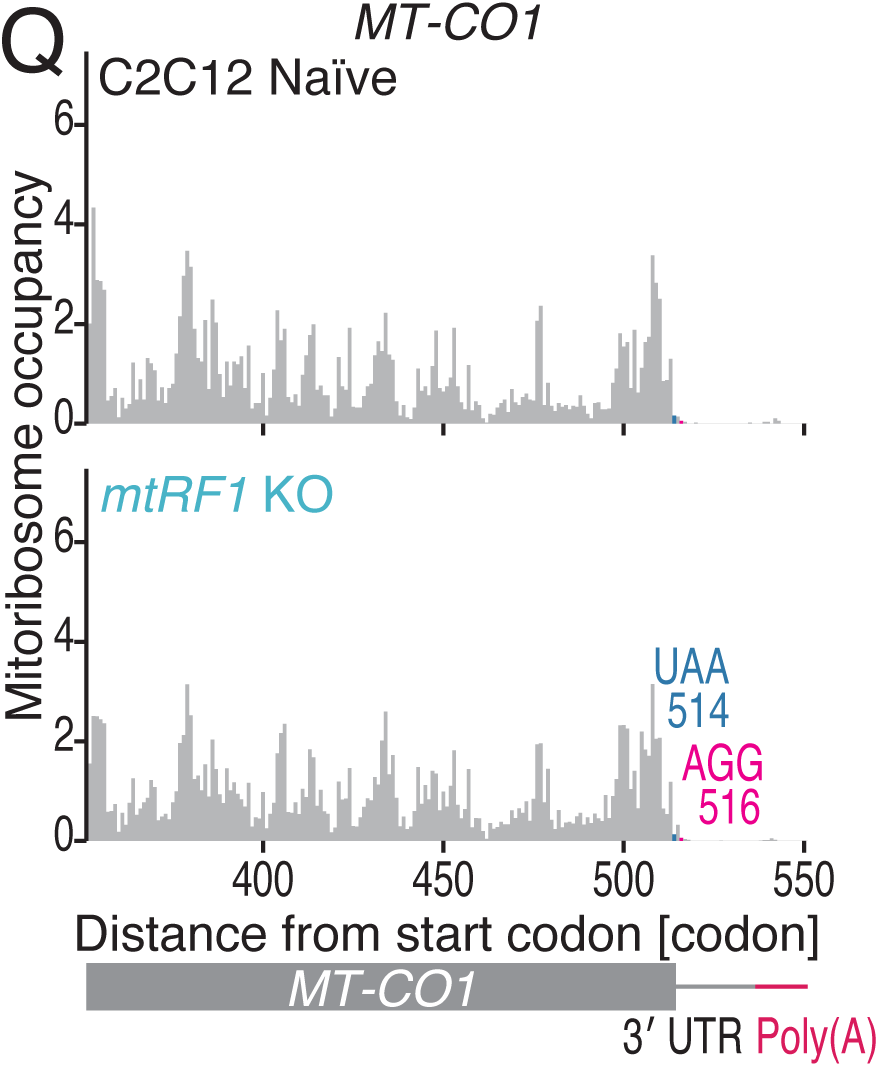
Characterization of MitoIP-Thor-Ribo-Seq and MitoIP-Thor-Disome-Seq data from *mtRF1* KO C2C12 cells, related to Figure 3. (A) Schematics showing the guide RNAs that were designed for CRISPR-Cas9-mediated gene knockout of mouse *mtRF1* and the deleted region in the isolated cell line. The genomic alterations in the isolated cell line are shown. (B) Genomic PCR of the *mtRF1* gene locus in the isolated cell line. The primer positions are depicted in A. (C) The loss of the mtRF1 protein in the isolated cell line was confirmed by Western blotting. The mitochondria-enriched fraction was subjected to immunoprecipitation with an anti-Tom22 antibody (MitoIP). Tom20 was used as a loading control. (D) Growth assay of naïve and *mtRF1* KO C2C12 cells grown under the indicated conditions. The means (points) and s.d.s (errors, n = 3) are shown. (E) Gel image of on-gel mito-FUNCAT experiments performed with the indicated cell lines. Proteins newly synthesized by mitoribosomes are labeled with L-homopropargyl glycine (HPG) and conjugated with infrared 800 (IR800) dye (left). The signal intensities along the lanes are shown (right). (F) Quantification of the on-gel mito-FUNCAT data in E. The means (bars), s.d.s (errors), and individual replicates (n = 3, points) are shown. CBB staining of total proteins was used for data normalization. (G) The fraction of read length of mitoribosome footprints in the indicated cells. Rep, replicate. (H) Polarity scores of the indicated ORFs in the indicated cells. The data points from the two replicates are shown individually. (I) Comparison of mitoribosome occupancy at the A-site (left) and P-site (right) codons in the indicated cells. (J) Metagene plots of mitoribosome footprints around canonical stop codons in the indicated cells. The A-site position of each read is displayed. The ORFs and stop codon sequences that were used in the analysis are shown. The *MT-ATP8* and *MT-ND4L* ORFs were excluded from the analysis since the 3′ ends of these ORFs overlap with the downstream *MT-ATP6* and *MT-ND4* ORFs in the polycistronic mRNAs. (K) The fraction of the read length of the mito-disome footprints in the indicated cells. Rep, replicate. (L) Comparison of mito-disome occupancy (for the leading mitoribosome) at the A-site (left) and P-site (right) codons in the indicated cells. (M) Metagene plots of mito-disome footprints around the canonical stop codons in the indicated cells. The A-site position (for the leading mitoribosome) of each read is displayed. The ORFs and stop codon sequences that were used in the analysis are shown. The *MT-ATP8* and *MT-ND4L* ORFs were excluded from the analysis since the 3′ ends of these ORFs overlap with the downstream *MT-ATP6* and *MT-ND4* ORFs in the polycistronic mRNAs. (N, O, and Q) Distribution of mitoribosome footprints in naïve and *mtRF1* KO C2C12 cells along the indicated transcripts. The A-site position of each read is displayed. The canonical and noncanonical stop codons are highlighted. (P) Schematics of human and mouse *MT-ATP8*/*MT-ATP6* polycistronic mRNAs. The reported ‒1 frameshift site in the *MT-ATP8* ORF and the secondary RNA structure in humans are shown. The orthogonal RNA structure (predicted) and codon/amino acid sequences in the ‒1 frame in mice are also shown.

**Figure S4.**
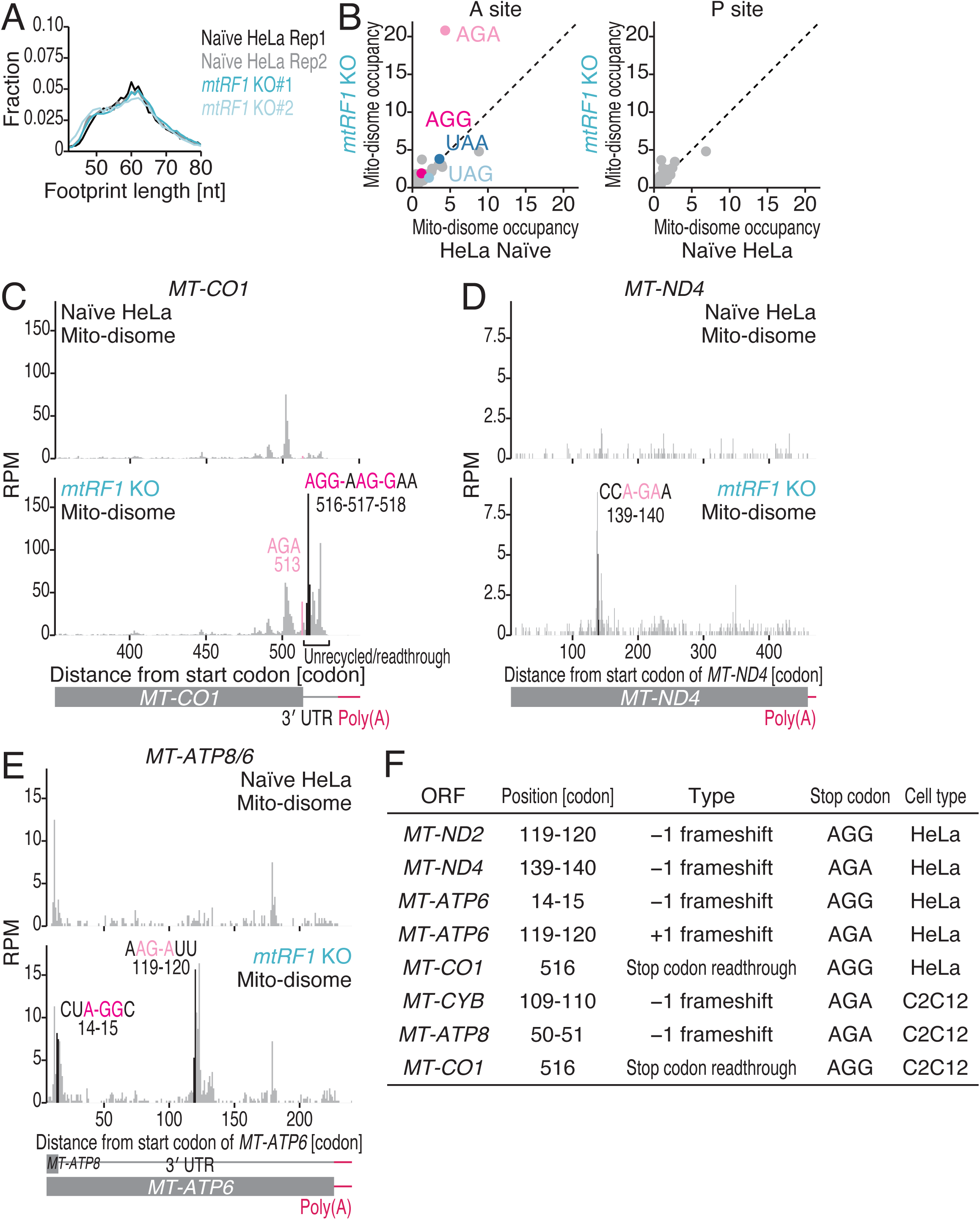
Characterization of MitoIP-Thor-Disome-Seq data from *mtRF1* KO HeLa cells, related to Figure 3. (A) The fraction of the read length of the mito-disome footprints in the indicated cells. Rep, replicate. (B) Comparison of mito-disome occupancy (for the leading mitoribosome) at the A-site (left) and P-site (right) codons in the indicated cells. (C-E) Distribution of mito-disome footprints in naïve and *mtRF1* KO HeLa cells along the indicated transcripts. The A-site position (for the leading mitoribosome) of each read is displayed. The in-frame and out-of-frame noncanonical stop codons with mito-disome peaks are highlighted. RPM, reads per million mapped reads. (F) Mito-disome peak sites found in *mtRF1* KO cells are listed with the ORF, position (the distance from start codons; *i.e.*, the start codon is counted as “0”), regulation type, stop codon sequence, and cell type.

**Figure S5.**
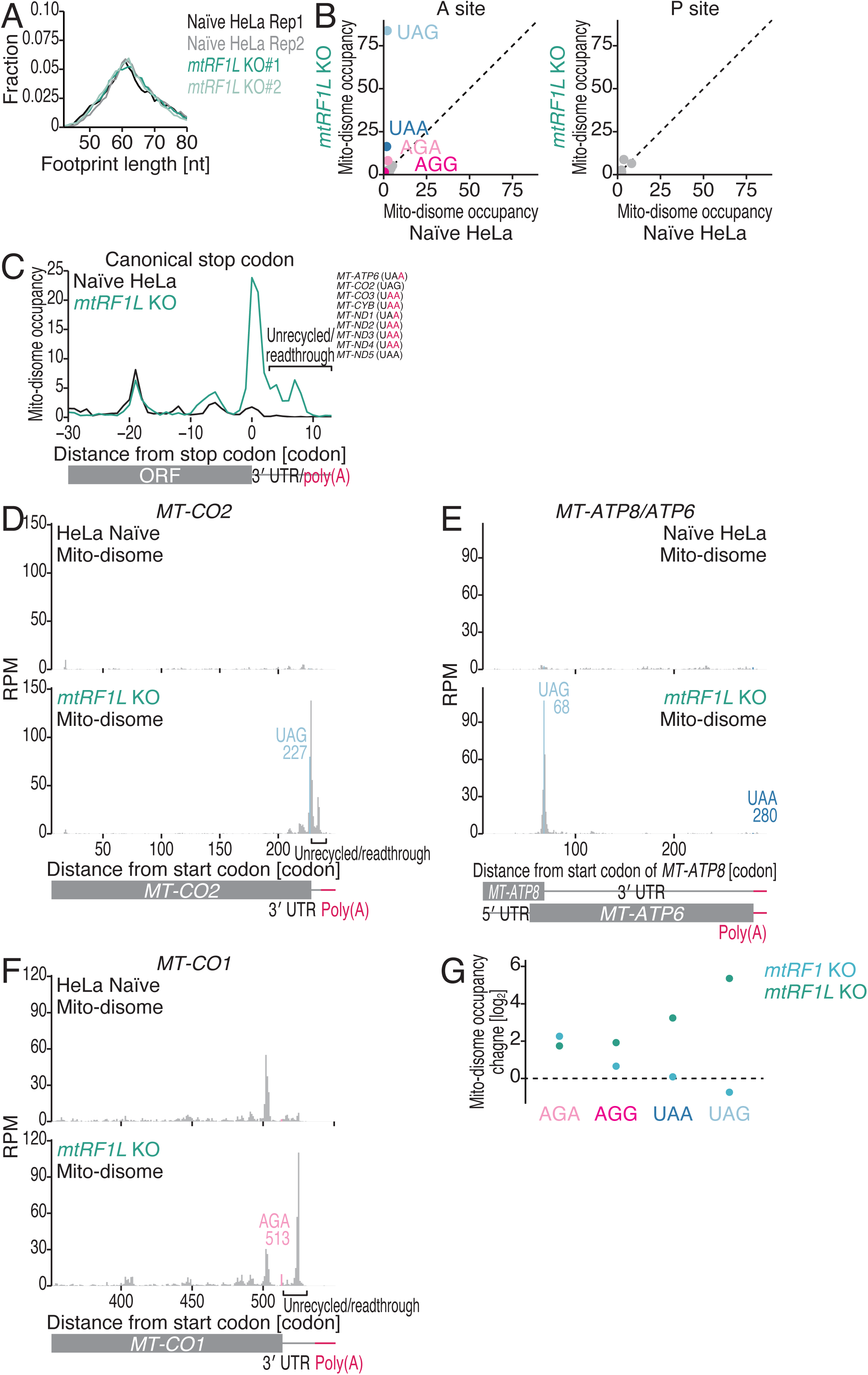
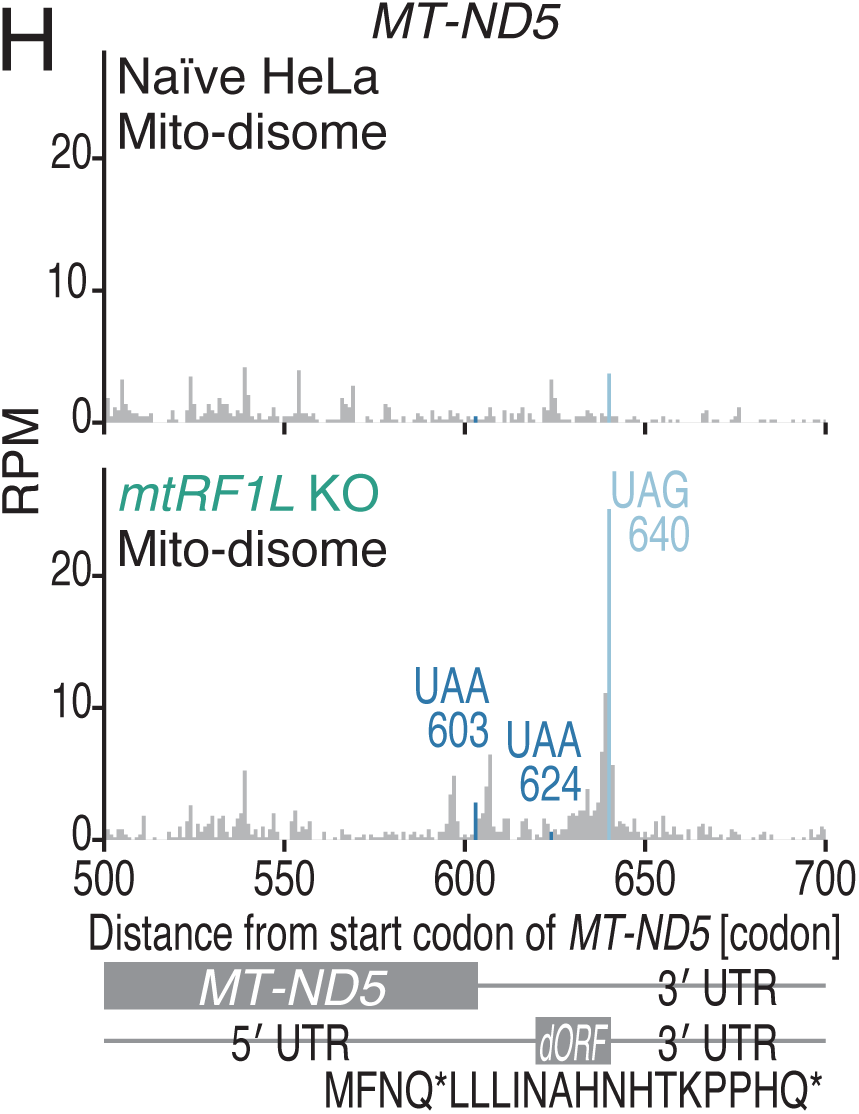
Characterization of MitoIP-Thor-Disome-Seq data from *mtRF1L* KO HeLa cells, related to Figure 3. (A) The fraction of the read length of the mito-disome footprints in the indicated cells. Rep, replicate. (B) Comparison of mito-disome occupancy (for the leading mitoribosome) at the A-site (left) and P-site (right) codons in the indicated cells. (C) Metagene plots of mito-disome footprints around the canonical stop codons in the indicated cells. The A-site position (for the leading mitoribosome) of each read is displayed. The ORFs and stop codon sequences that were used in the analysis are shown. The *MT-ATP8* and *MT-ND4L* ORFs were excluded from the analysis since the 3′ ends of these ORFs overlap with the downstream *MT-ATP6* and *MT-ND4* ORFs in the polycistronic mRNAs. (D, E, F, and H) Distribution of mito-disome footprints in naïve and *mtRF1* KO HeLa cells along the indicated transcripts. The A-site position (for the leading mitoribosome) of each read is displayed. The in-frame canonical and noncanonical stop codons with mito-disome peaks are highlighted. RPM, reads per million mapped reads. (G) Changes in mito-disome occupancy (for the leading mitoribosome) at stop codons in the indicated cells.

**Figure S6.**
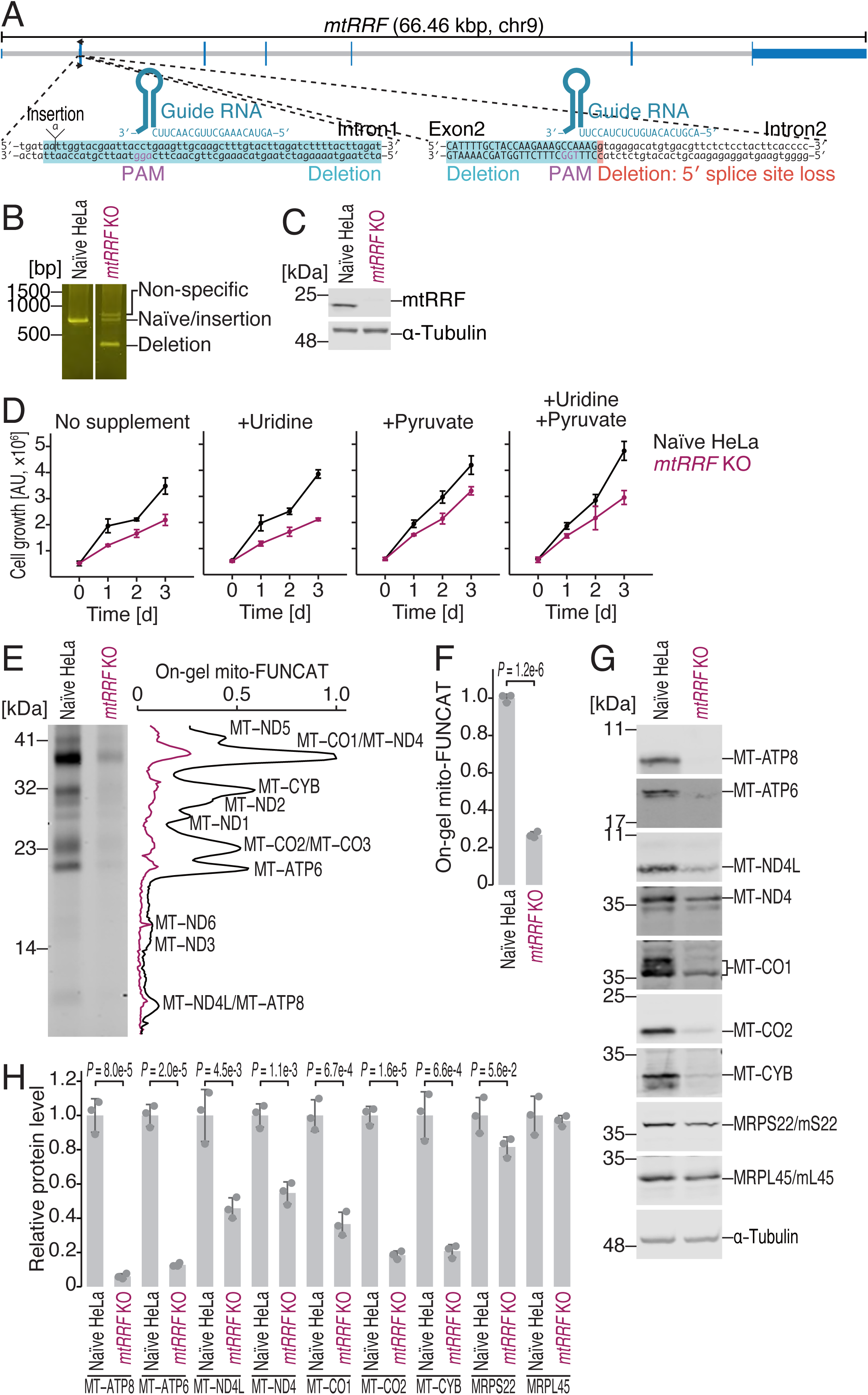
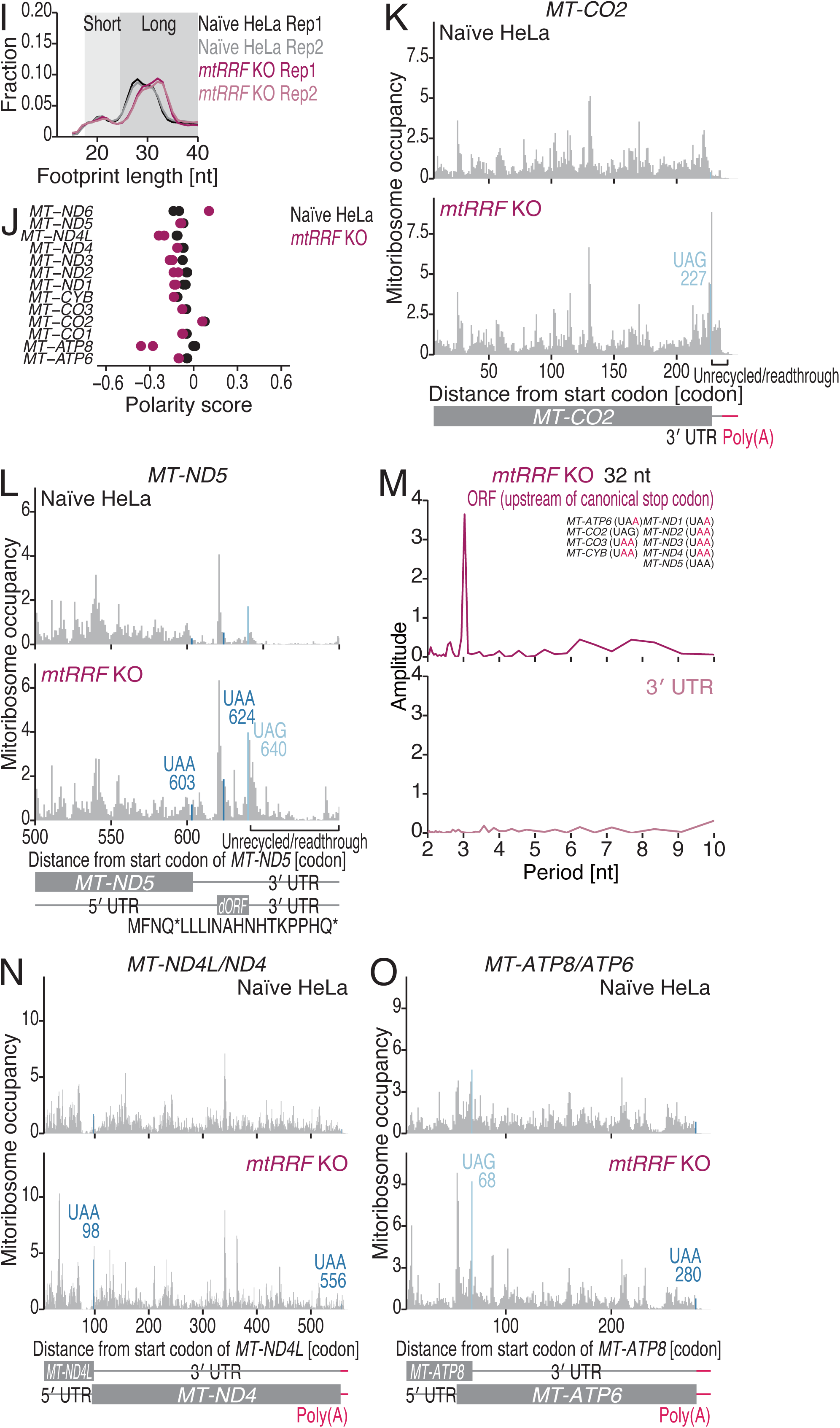
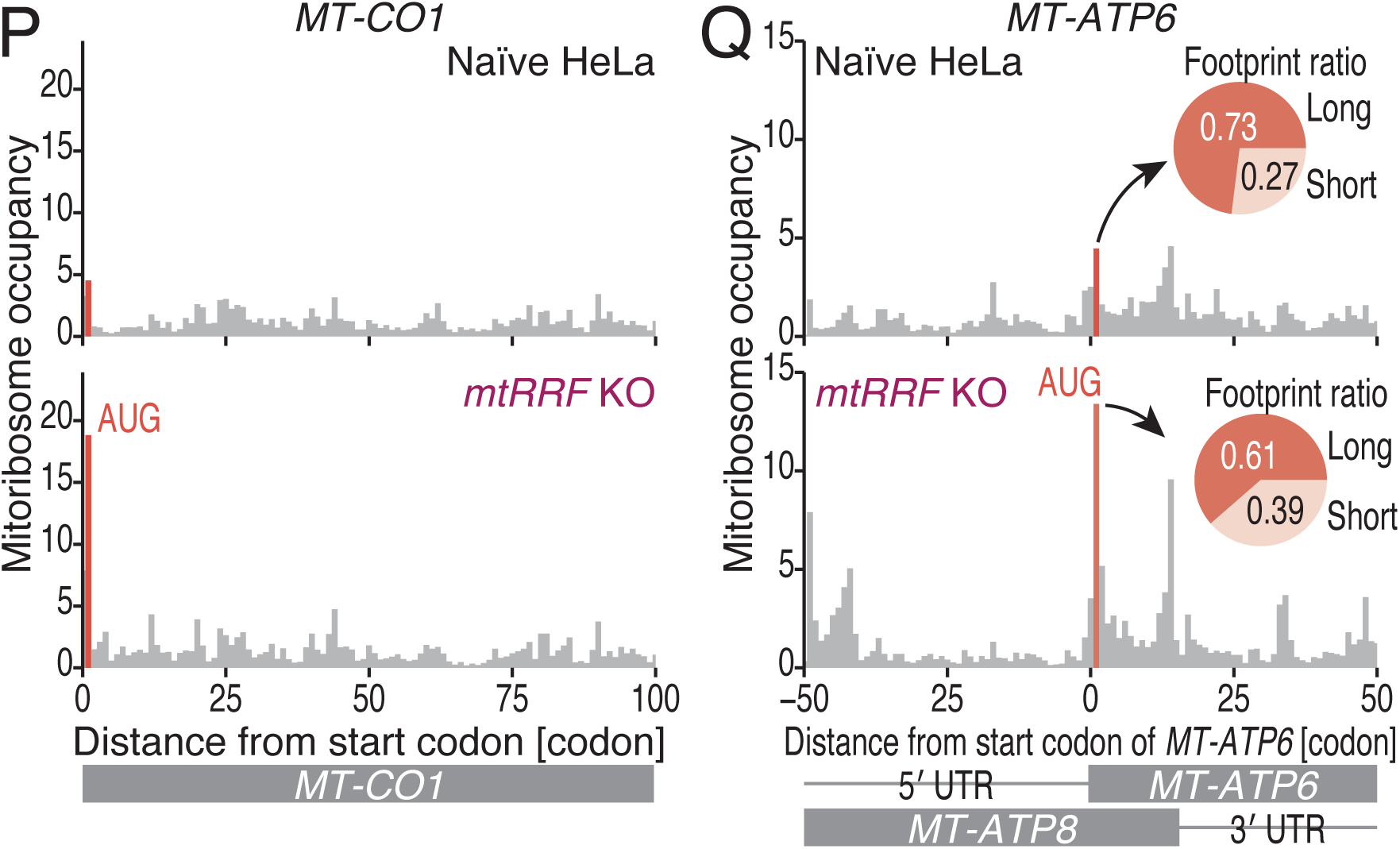
Characterization of MitoIP-Thor-Ribo-Seq data from *mtRRF* KO HeLa cells, related to Figure 4. (A) Schematics showing the guide RNAs that were designed for CRISPR-Cas9-mediated gene knockout of human *mtRRF* and the deleted region in the isolated cell line. The genomic alterations in the isolated cell line are shown. (B) Genomic PCR of the *mtRRF* gene locus in the isolated cell line. The primer positions are depicted in A. (C) The loss of the mtRRF protein in the isolated cell line was confirmed by Western blotting. Total lysate was used. α-Tubulin was used as a loading control. (D) Growth assay of naïve and *mtRRF* KO HeLa cells grown under the indicated conditions. The means (points) and s.d.s (errors, n = 3) are shown. (E) Gel image of on-gel mito-FUNCAT experiments performed with the indicated cell lines. Proteins newly synthesized by mitoribosomes are labeled with L-homopropargyl glycine (HPG) and conjugated with infrared 800 (IR800) dye (left). The signal intensities along the lanes are shown (right). (F) Quantification of the on-gel mito-FUNCAT data in E. The means (bars), s.d.s (errors), and individual replicates (n = 3, points) are shown. CBB staining of total proteins was used for data normalization. The p values were calculated with Student’s t test (two-tailed). (G) Western blotting for the indicated proteins in the indicated cell lines. α-Tubulin was used as a loading control. (H) Quantification of the on-gel mito-FUNCAT data in G. The means (bars), s.d.s (errors), and individual replicates (n = 3, points) are shown. α-Tubulin was used for data normalization. The p values were calculated with Student’s t test (two-tailed). (I) The fraction of read length of mitoribosome footprints in the indicated cells. Rep, replicate. (J) Polarity scores of the indicated ORFs in the indicated cells. The data points from the two replicates are shown individually. (K, L, and N-Q) Distribution of mitoribosome footprints in naïve and *mtRRF* KO HeLa cells along the indicated transcripts. The A-site position of each read is displayed. For K, L, N, and O, the canonical stop codon is highlighted. For N and O, the start codon is highlighted. In Q, the inset pie chart represents the fractions of long footprints (25-40 nt) and short footprints (18-24 nt) at the start codon of *MT-ATP6*. (M) Discrete Fourier transform of mitoribosome footprints mapped to the 100-nt region upstream and downstream of canonical stop codons. The ORFs and stop codon sequences that were used in the analysis are shown.

**Figure S7.**
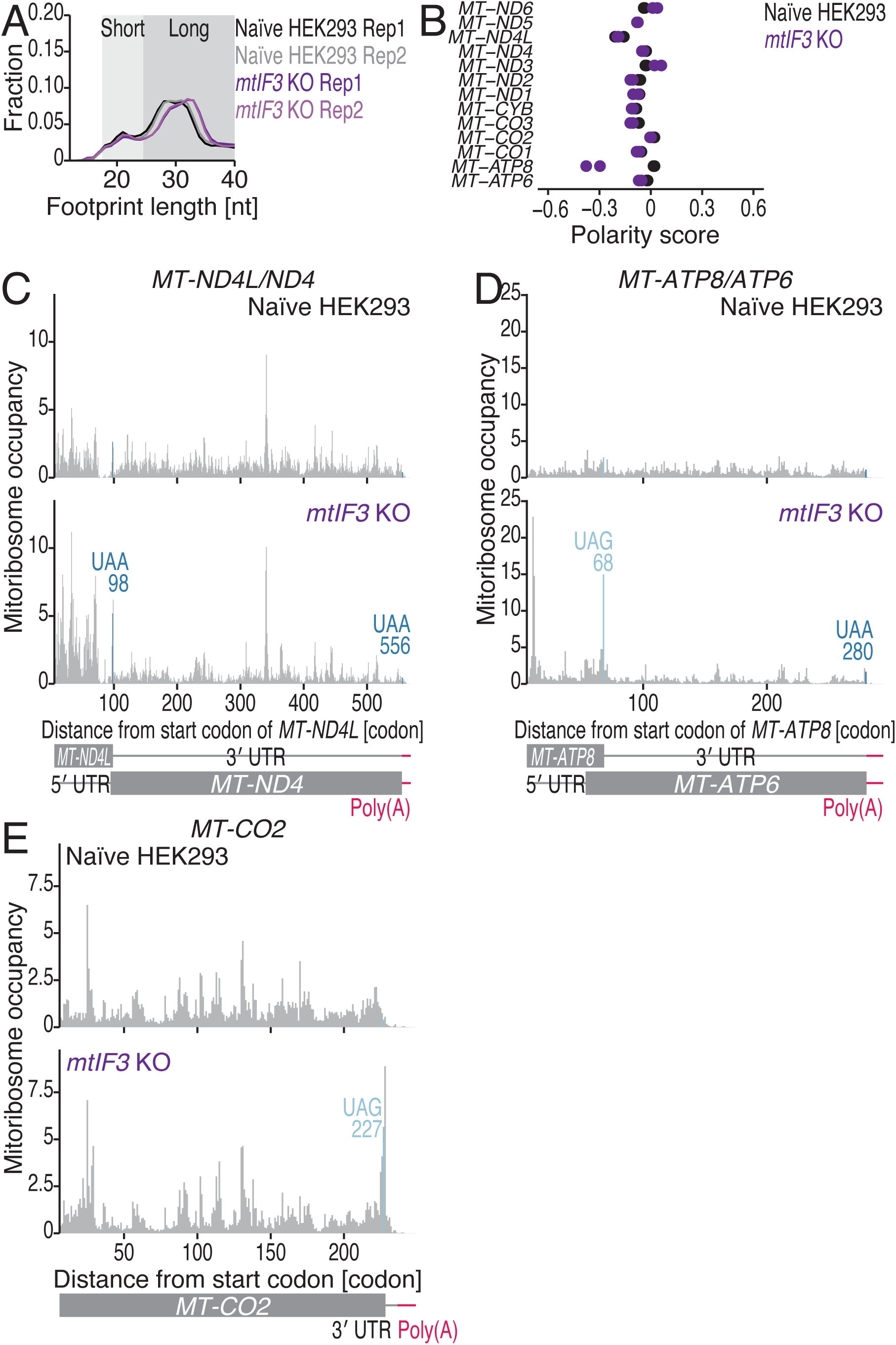
Characterization of MitoIP-Thor-Ribo-Seq data from *mtIF3* KO HeLa cells, related to Figure 5. (A) The fraction of read length of mitoribosome footprints in the indicated cells. Rep, replicate. (B) Polarity scores of the indicated ORFs in the indicated cells. The data points from the two replicates are shown individually. (C-E) Distribution of mitoribosome footprints in naïve and *mtRRF* KO HeLa cells along the indicated transcripts. The A-site position of each read is displayed. The canonical stop codon is highlighted.

**Figure S8.**
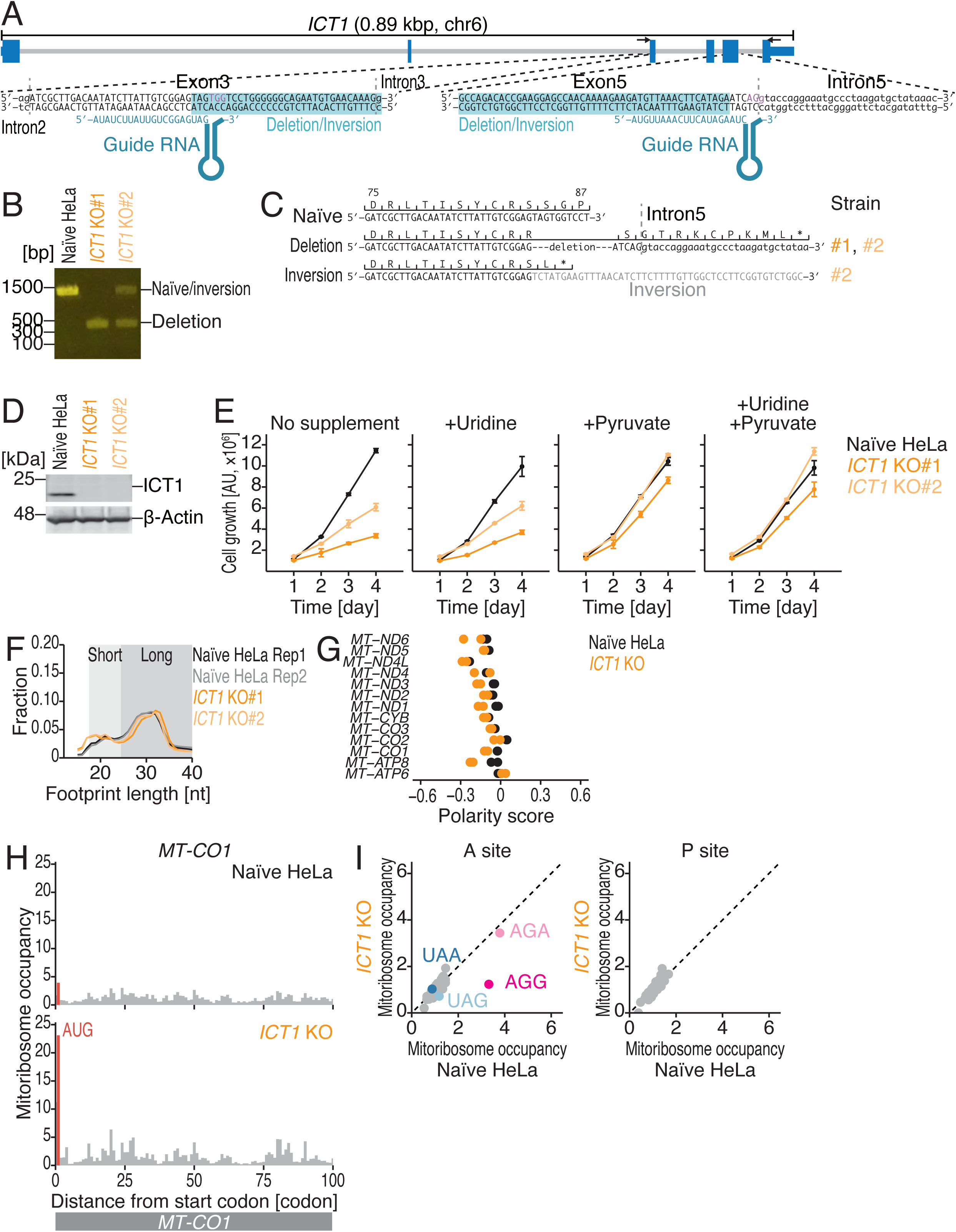
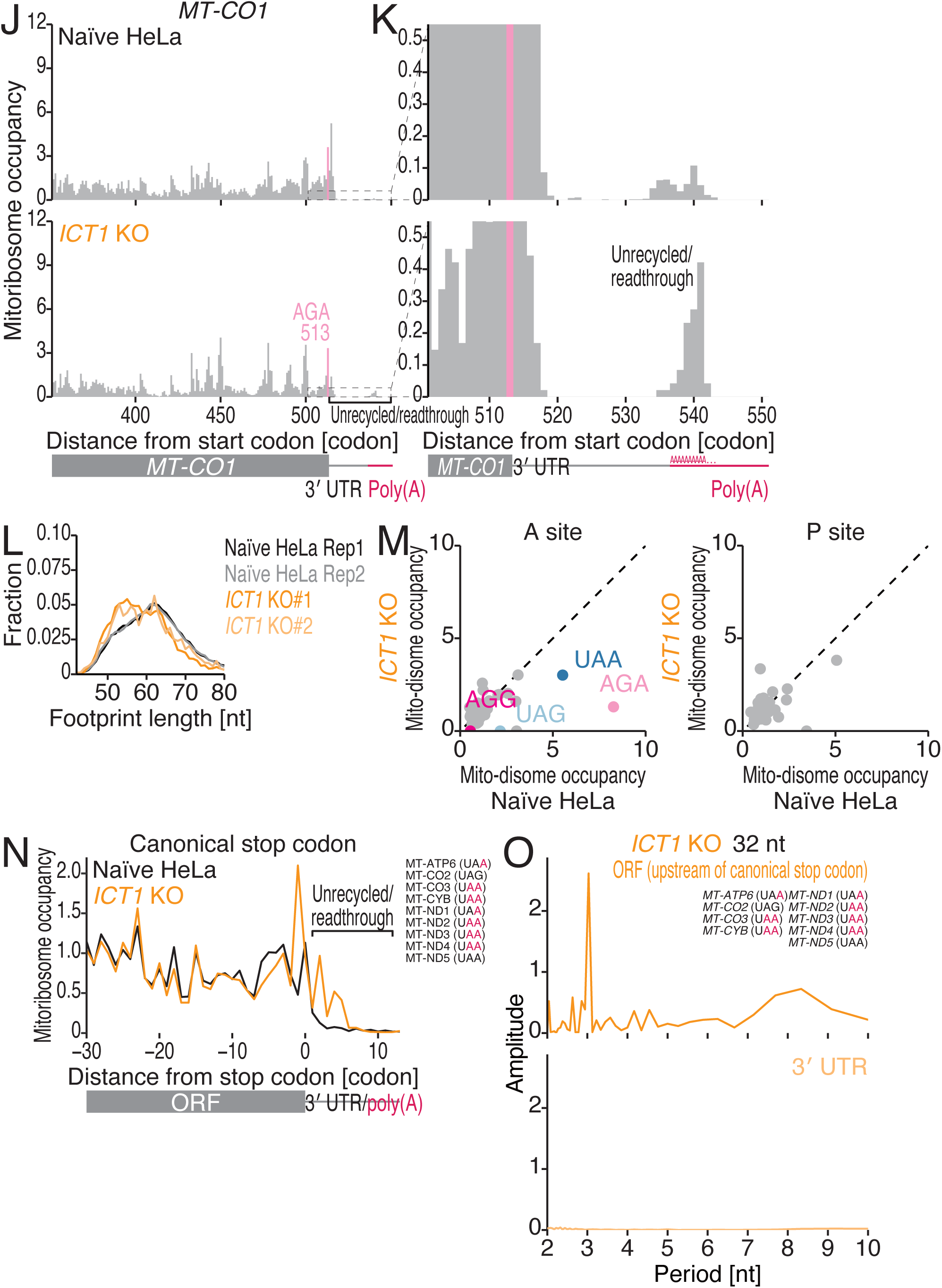
Characterization of MitoIP-Thor-Ribo-Seq and MitoIP-Thor-Disome-Seq data from *ICT1* KO HeLa cells, related to Figure 6. (A) Schematics showing the guide RNAs that were designed for CRISPR-Cas9-mediated gene knockout of human *ICT1* and the deleted region in the isolated cell lines. (B) Genomic PCR of the *ICT1* gene locus in the isolated cell lines. The primer positions are depicted in A. (C) Schematic of the genomic alterations in the isolated cell lines. (D) The loss of ICT1 protein in the isolated cell lines was confirmed by Western blotting. Total lysate was used. β-Actin was used as a loading control. (E) Growth assay of naïve and *ICT1* KO HeLa cells grown under the indicated conditions. The means (points) and s.d.s (errors, n = 3) are shown. (F) The fraction of read length of mitoribosome footprints in the indicated cells. Rep, replicate. (G) Polarity scores of the indicated ORFs in the indicated cells. The data points from the two replicates are shown individually. (H, J, and K) Distribution of mitoribosome footprints in naïve and *ICT1* KO HeLa cells along the indicated transcripts. The A-site position of each read is displayed. In H, the start codon is highlighted. In J and K, the noncanonical stop codons are highlighted. K represents the magnified view of J downstream of the stop codon. (I) Comparison of mitoribosome occupancy at the A-site (left) and P-site (right) codons in the indicated cells. (L) The fraction of the read length of the mito-disome footprints in the indicated cells. Rep, replicate. (M) Comparison of mito-disome occupancy (for the leading mitoribosome) at the A-site (left) and P-site (right) codons in the indicated cells. (N) Metagene plots of mitoribosome footprints around canonical stop codons in the indicated cells. The A-site position of each read is displayed. The ORFs and stop codon sequences that were used in the analysis are shown. The *MT-ATP8* and *MT-ND4L* ORFs were excluded from the analysis since the 3′ ends of these ORFs overlap with the downstream *MT-ATP6* and *MT-ND4* ORFs in the polycistronic mRNAs. (O) Discrete Fourier transform of mitoribosome footprints mapped to the 100-nt region upstream and downstream of canonical stop codons. The ORFs and stop codon sequences that were used in the analysis are shown.

**Figure S9.**
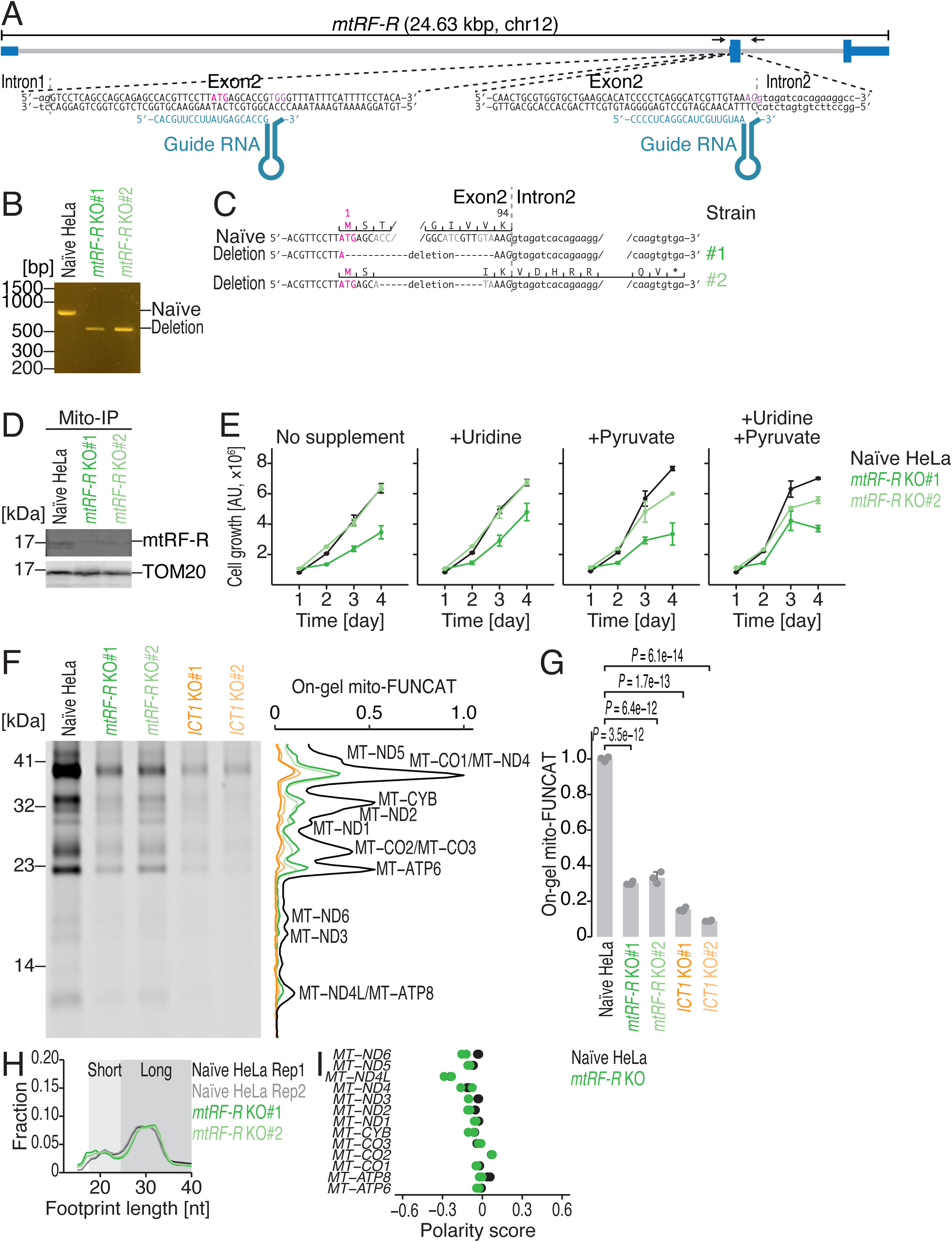
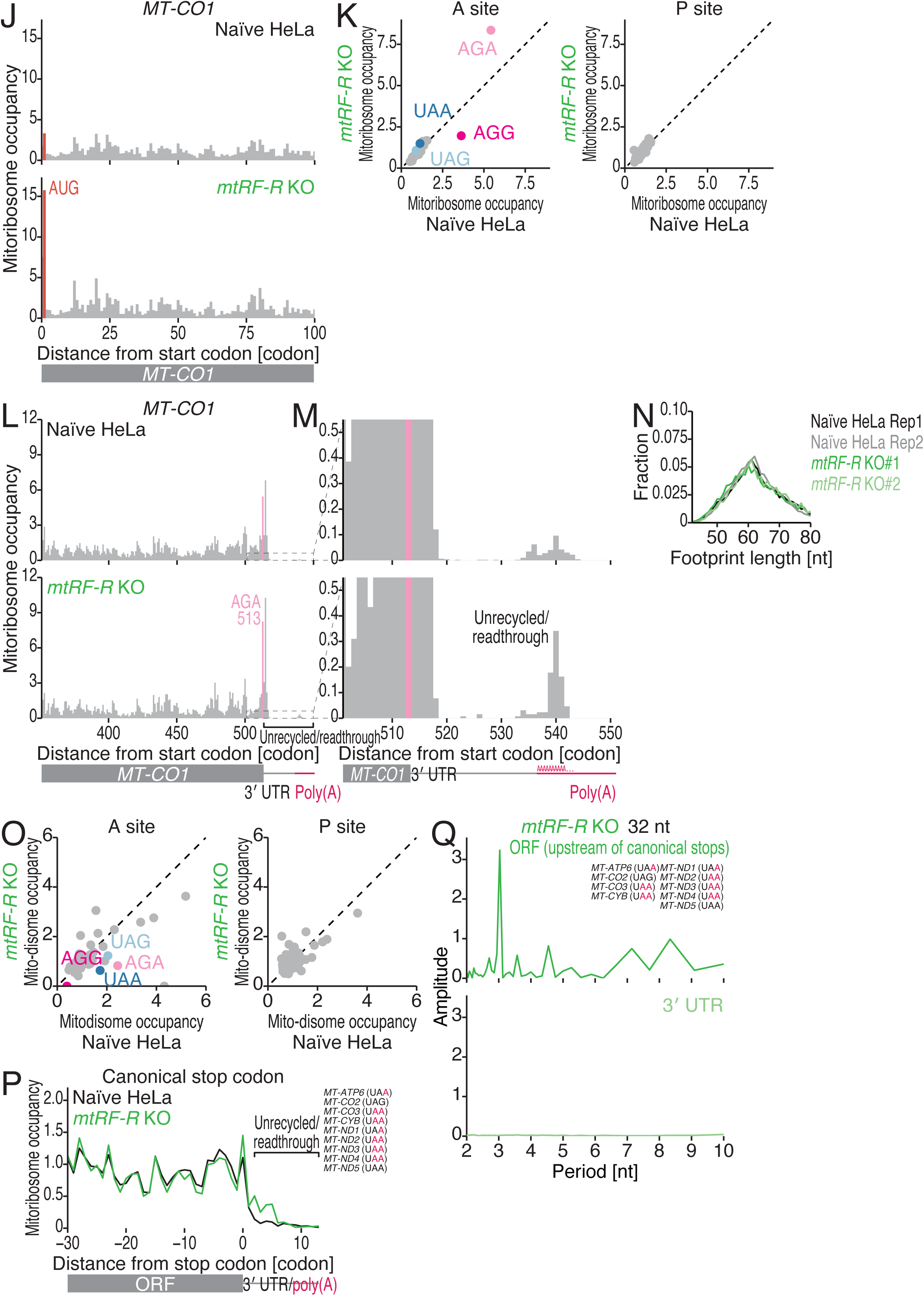
Characterization of MitoIP-Thor-Ribo-Seq and MitoIP-Thor-Disome-Seq data from *mtRF-R* KO HeLa cells, related to Figure 6. (A) Schematics showing the guide RNAs that were designed for CRISPR-Cas9-mediated gene knockout of human *mtRF-R* and the deleted region in the isolated cell lines. (B) Genomic PCR of the *mtRF-R* gene locus in the isolated cell lines. The primer positions are depicted in A. (C) Schematic of the genomic alterations in the isolated cell lines. (D) The loss of mtRF-R protein in the isolated cell lines was confirmed by Western blotting. The mitochondria-enriched fraction was subjected to immunoprecipitation with an anti-TOM22 antibody (MitoIP). TOM20 was used as a loading control. (E) Growth assay of naïve and *mtRF-R* KO HeLa cells grown under the indicated conditions. The means (points) and s.d.s (errors, n = 3) are shown. (F) Gel image for on-gel mito-FUNCAT experiments in the indicated cell lines. Proteins newly synthesized by mitoribosomes are labeled with L-homopropargyl glycine (HPG) and conjugated with infrared 800 (IR800) dye (left). The signal intensities along the lanes are shown (right). We note that the data for the naïve samples are the same as those in Figure S1N. (G) Quantification of the on-gel mito-FUNCAT data in F. The means (bars), s.d.s (errors), and individual replicates (n = 3, points) are shown. CBB staining of total proteins was used for data normalization. The p values were calculated with the Tukey‒Kramer test (two-tailed). We note that naïve samples are the same data used for Figure S1O. (H) The fraction of read length of mitoribosome footprints in the indicated cells. Rep, replicate. (I) Polarity scores of the indicated ORFs in the indicated cells. The data points from the two replicates are shown individually. (J, L, and M) Distribution of mitoribosome footprints in naïve and *ICT1* KO HeLa cells along the indicated transcripts. The A-site position of each read is displayed. In J, the start codon is highlighted. In L-M, the noncanonical stop codons are highlighted. M represents the magnified view of L downstream of the stop codon. (K) Comparison of mitoribosome occupancy at the A-site (left) and P-site (right) codons in the indicated cells. (N) The fraction of the read length of the mito-disome footprints in the indicated cells. Rep, replicate. (O) Comparison of mito-disome occupancy (for the leading mitoribosome) at the A-site (left) and P-site (right) codons in the indicated cells. (P) Metagene plots of mitoribosome footprints around canonical stop codons in the indicated cells. The A-site position of each read is displayed. The ORFs and stop codon sequences that were used in the analysis are shown. The *MT-ATP8* and *MT-ND4L* ORFs were excluded from the analysis since the 3′ ends of these ORFs overlap with the downstream *MT-ATP6* and *MT-ND4* ORFs in the polycistronic mRNAs. (Q) Discrete Fourier transform of mitoribosome footprints mapped to the 100-nt region upstream and downstream of canonical stop codons. The ORFs and stop codon sequences that were used in the analysis are shown.

**Figure S10.**
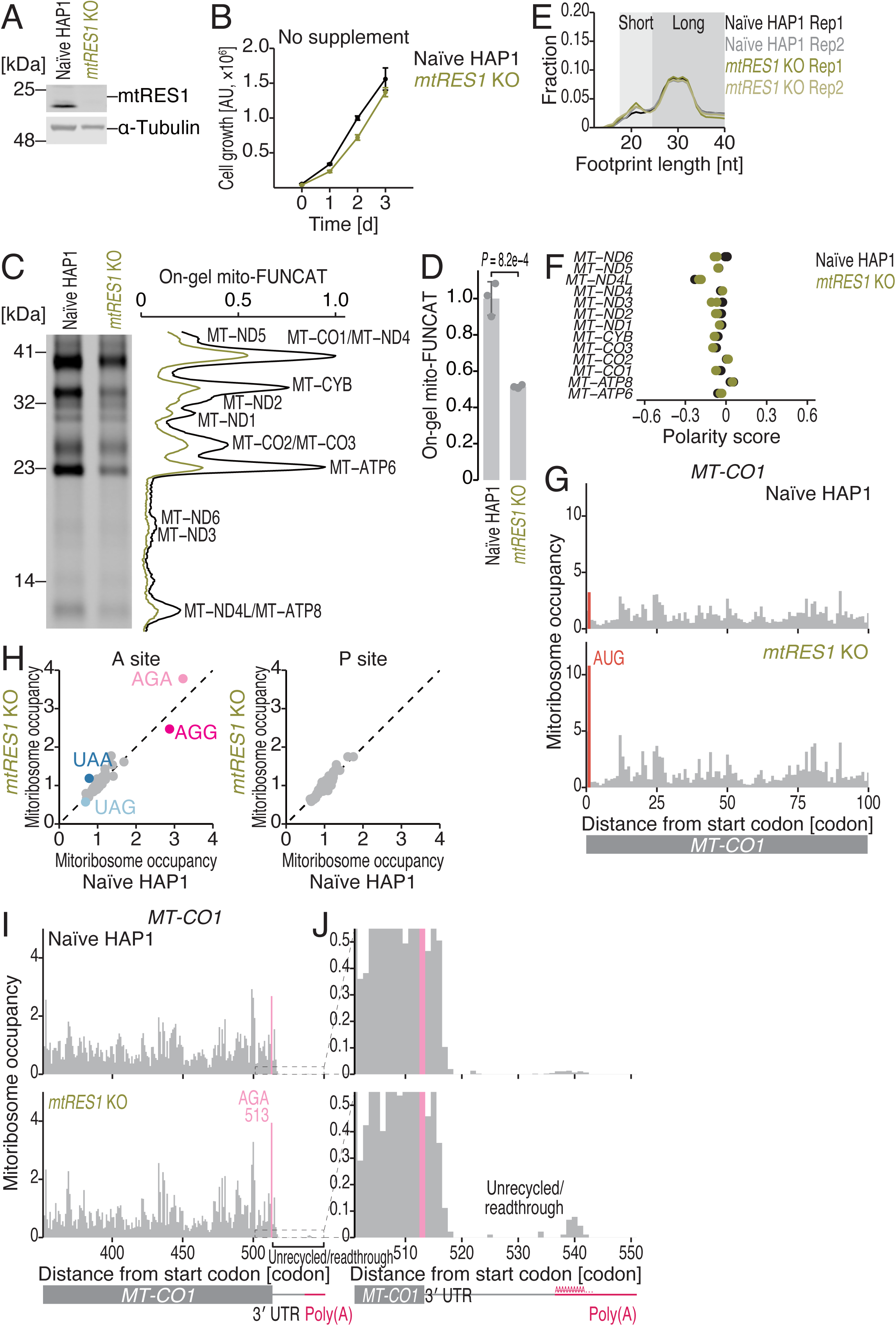
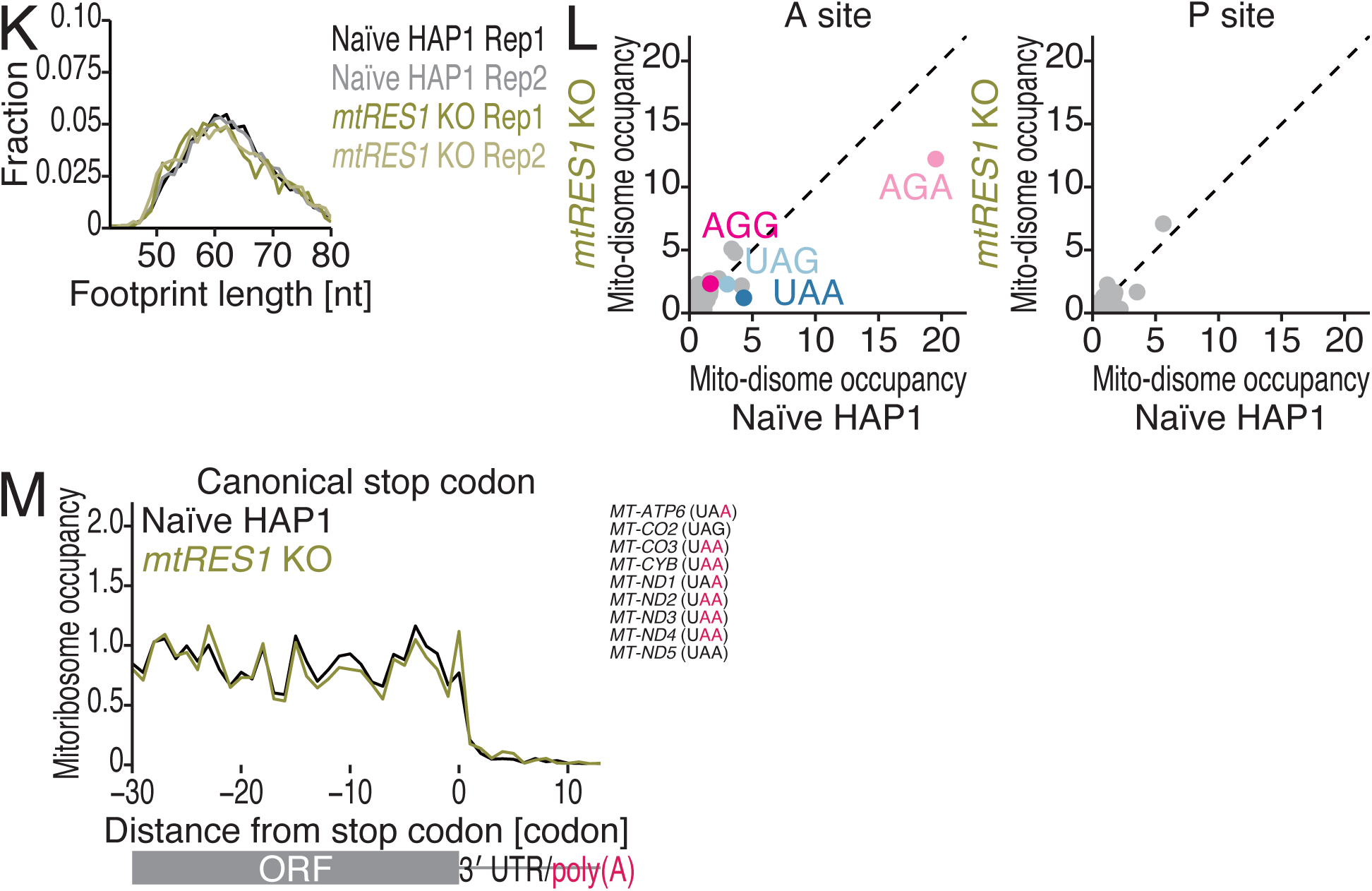
Characterization of MitoIP-Thor-Ribo-Seq and MitoIP-Thor-Disome-Seq data from *mtRES1* KO HAP1 cells, related to Figure 6. (A) The loss of the mtRES1 protein in the cell line was confirmed by Western blotting. Total lysate was used. α-Tubulin was used as a loading control. (B) Growth assay of naïve and *mtRES1-*KO HAP1 cells grown under the indicated conditions. The means (points) and s.d.s (errors, n = 3) are shown. (C) Gel image of on-gel mito-FUNCAT experiments performed with the indicated cell lines. Proteins newly synthesized by mitoribosomes are labeled with L-homopropargyl glycine (HPG) and conjugated with infrared 800 (IR800) dye (left). The signal intensities along the lanes are shown (right). (D) Quantification of the on-gel mito-FUNCAT data in C. The means (bars), s.d.s (errors), and individual replicates (n = 3, points) are shown. CBB staining of total proteins was used for data normalization. The p values were calculated with Student’s t test (two-tailed). (E) The fraction of read length of mitoribosome footprints in the indicated cells. Rep, replicate. (F) Polarity scores of the indicated ORFs in the indicated cells. The data points from the two replicates are shown individually. (G, I, and J) Distribution of mitoribosome footprints in naïve and *ICT1* KO HeLa cells along the indicated transcripts. The A-site position of each read is displayed. In G, the start codon is highlighted. In I and J, the noncanonical stop codons are highlighted. J represents the magnified view of I downstream of the stop codon. (H) Comparison of mitoribosome occupancy at the A-site (left) and P-site (right) codons in the indicated cells. (K) The fraction of the read length of the mito-disome footprints in the indicated cells. Rep, replicate. (L) Comparison of mito-disome occupancy (for the leading mitoribosome) at the A-site (left) and P-site (right) codons in the indicated cells. (M) Metagene plots of mitoribosome footprints around canonical stop codons in the indicated cells. The A-site position of each read is displayed. The ORFs and stop codon sequences that were used in the analysis are shown. The *MT-ATP8* and *MT-ND4L* ORFs were excluded from the analysis since the 3′ ends of these ORFs overlap with the downstream *MT-ATP6* and *MT-ND4* ORFs in the polycistronic mRNAs.

**Figure S11.**
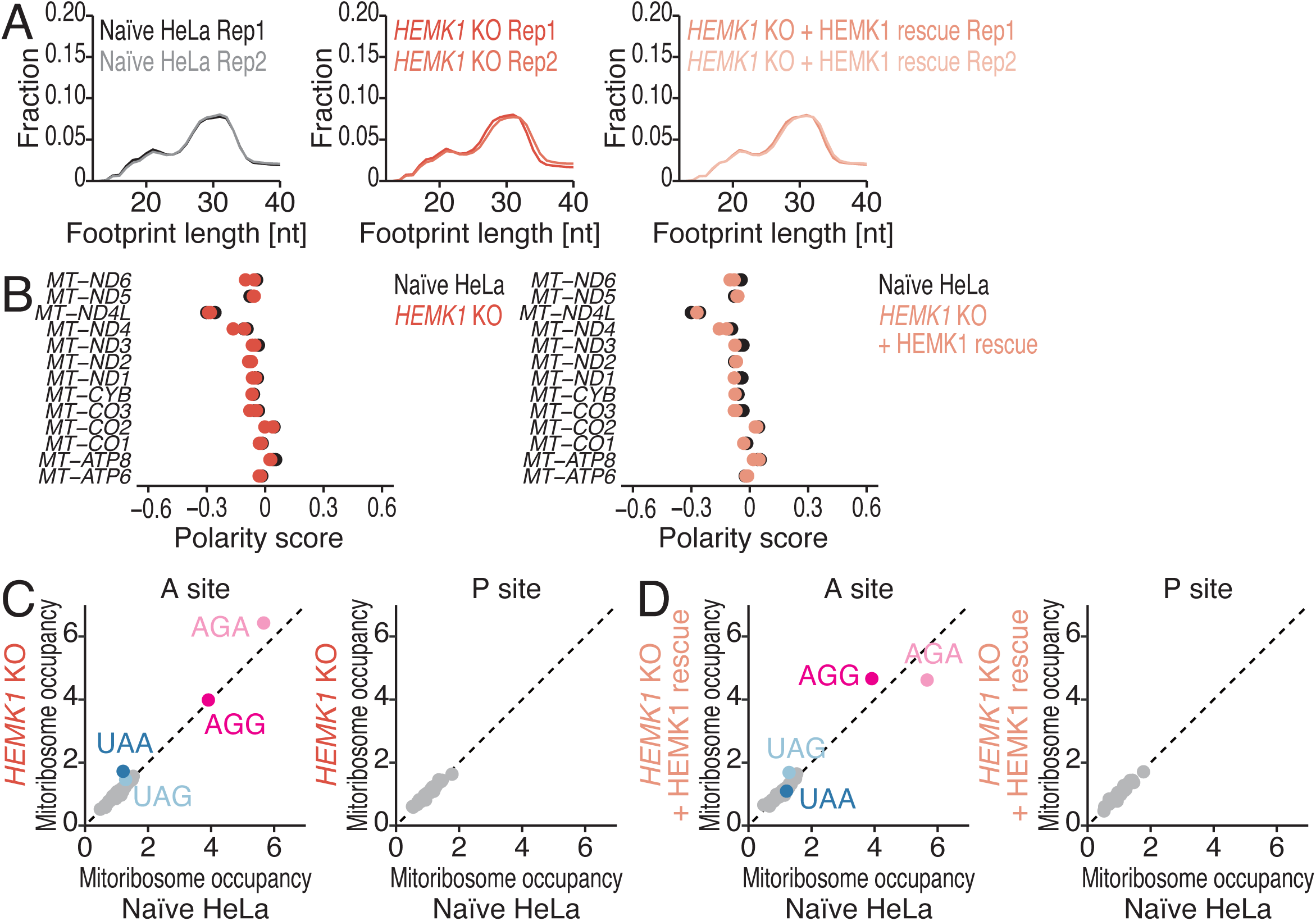
Characterization of MitoIP-Thor-Ribo-Seq and MitoIP-Thor-Disome-Seq data from *HEMK1* KO HeLa cells, related to Figure 7. (A) The fraction of read length of mitoribosome footprints in the indicated cells. Rep, replicate. (B) Polarity scores of the indicated ORFs in the indicated cells. The data points from the two replicates are shown individually. (C and D) Comparison of mitoribosome occupancy at the A-site (left) and P-site (right) codons in the indicated cells.

**Table S1. Summary of MitoIP-Thor-Ribo-Seq and MitoIP-Thor-Disome-Seq data obtained in this study, related to Figures 1-7**.

For each dataset, the library name, culture condition, sample name, cell line name, library type, rRNA depletion method, mitoribosome footprint length analyzed, A-site offset for each mitoribosome footprint length, cytoribosome footprint length analyzed, A-site offset for each cytoribosome footprint length, accession number, and reference are listed. Rep, replicate.

